# The therapeutic and diagnostic potential of amyloid β oligomers (AβOs) selective antibodies to treat Alzheimer’s disease

**DOI:** 10.1101/2021.09.20.460817

**Authors:** Kirsten L. Viola, Maira A. Bicca, Adrian M. Bebenek, Daniel L. Kranz, Vikas Nandwana, Emily A. Waters, Chad R. Haney, Maxwell Lee, Abhay Gupta, Zach Brahmbhatt, Weijian Huang, Ting-Tung Chang, Anderson Peck, Clarissa Valdez, Vinayak P. Dravid, William L. Klein

## Abstract

Improvements have been made in the diagnosis of Alzheimer’s disease (AD), manifesting mostly in the development of *in vivo* imaging methods that allow for the detection of pathological changes in AD by MRI and PET scans. Many of these imaging methods, however, use agents that probe amyloid fibrils and plaques - species that do not correlate well with disease progression and are not present at the earliest stages of the disease. Amyloid β oligomers (AβOs), rather, are now widely accepted as the Aβ species most germane to AD onset and progression. Here we report evidence further supporting the role of AβOs as pathological instigators of AD and introduce promising anti-AβO diagnostic probes capable of distinguishing the 5xFAD mouse model from wild type mice by PET and MRI. In a developmental study, Aβ oligomers in 5xFAD mice were found to appear at 3 months of age, just prior to the onset of memory dysfunction, and spread as memory worsened. The increase of AβOs is prominent in the subiculum and correlates with concomitant development of reactive astrocytosis. The impact of these AβOs on memory is in harmony with findings that intraventricular injection of synthetic AβOs into wild type mice induced hippocampal dependent memory dysfunction within 24 hours. Compelling support for the conclusion that endogenous AβOs cause memory loss was found in experiments showing that intranasal inoculation of AβO-selective antibodies into 5xFAD mice completely restored memory function, measured 30-40 days post-inoculation. These antibodies, which were modified to give MRI and PET imaging probes, were able to distinguish 5xFAD mice from wild type littermates. These results provide strong support for the role of AβOs in instigating memory loss and salient AD neuropathology, and they demonstrate that AβO selective antibodies have potential both for therapeutics and for diagnostics.

## Introduction

### General Alzheimer’s disease

More than 6 million Americans are currently living with Alzheimer’s disease (AD), and Alzheimer’s-related deaths have increased 145% from 2000 to 2019 (2021). The financial burden is even more staggering - Alzheimer’s and other dementias have cost the US more than $600 billion in medical expenses and unpaid care in 2021 (2021). Despite the great personal and economic burden, progress toward developing effective diagnostics and therapeutics remains slow. Aduhelm^®^ (also known as Aducanumab) was recently approved as a treatment for AD (Investor Relations, 2021), the first in more than a decade, but it still focuses on Aβ elimination rather than specific AβO targets. As AD burden is expected to increase drastically with the aging population, improved diagnostics and therapeutics are more urgent now than ever.

### AβOs as a biomarker for early Alzheimer’s disease

The primary pathological hallmarks of Alzheimer’s disease are extracellular amyloid plaques and intraneuronal tangles of hyperphosphorylated tau (Masters et al., 1985). It is well known, however, that amyloid plaques do not correlate well with cognitive decline in AD (Terry et al., 1991; Hsia et al., 1999; Lee et al., 2004) and are not present in the earliest stages of the disease (Nyborg et al., 2013). Research from the previous two decades strongly indicates that soluble amyloid beta oligomers (AβOs), not plaques, are the more appropriate amyloid beta species to target in AD (Ashe, 2020; Hampel et al., 2021).

AβOs are potent neurotoxins that show AD-dependent accumulation in the brain of AD patients (Gong et al., 2003; Kayed et al., 2003; Lacor et al., 2004) and transgenic (Tg) rodent AD models (Chang et al., 2003; Lesne et al., 2006; Ohno et al., 2006). For reviews of other perspectives regarding AD molecular etiology, see (Braak and Del Tredici, 2011; Robakis, 2011; Lasagna-Reeves et al., 2012). AβOs begin to accumulate early in AD, decades prior to symptoms, and are widely held to be the neurotoxic instigators of AD (Rodgers, 2005; Gandy et al., 2010; Schnabel, 2011; Mucke and Selkoe, 2012). AβOs have been shown to exert their toxic effects by instigating failure of synaptic plasticity and memory (Lambert et al., 1998; Wang et al., 2002; Lesne et al., 2006; Townsend et al., 2006). Recently, soluble cortical extracts were examined by ELISA and showed that the ratio of AβO levels to plaque density fully distinguished demented from non-demented patients (Esparza et al., 2013); simply put, those with high AβO to plaque ratios were demented and low AβO to plaque ratios were not.

### The 5xFAD mouse model

The 5xFAD transgenic mouse is an increasingly used AD model that harbors gene mutations in amyloid β protein precursor (AβPP) (K670N/M671L + I716V + V717I) and presenilins (PS1/2) (M146L + L286V) (Oakley et al., 2006). These mutations are known to increase production of Aβ42, characteristic of familial AD, and exhibit expedited plaque development compared to other transgenic mice (Oakley et al., 2006). The Mutant Mouse Resource Research Center (MMRRC) found that Aβ accumulation occurred at different rates, depending on the breeding background, with mice bred on a B6SJL background developing pathology at a significantly more rapid rate (unpublished, available at <u>MMRRC 5xFAD strain data</u>) than those bred on a C57 background. The 5xFAD mouse model is well characterized for memory impairments (Oakley et al., 2006; Kimura and Ohno, 2009; Ohno, 2009; Girard et al., 2013; Girard et al., 2014; Zhang et al., 2021a), neuron loss (Jawhar et al., 2012; Oblak et al., 2021), and Aβ plaque accumulation (Devi et al., 2010; Jawhar et al., 2012; Ashe, 2020; Zhang et al., 2021a). Comprehensive studies on the 5xFAD model have also looked at cholesterol and glucose levels (Oblak et al., 2021), activity levels (Oblak et al., 2021), neuroinflammation-related protein levels (Ou-Yang and Van Nostrand, 2013; Oblak et al., 2021), tau phosphorylation (Shao et al., 2011; Kanno et al., 2014), and visual acuity (Zhang et al., 2021a).

### Alzheimer’s disease diagnostics

Recommended tests (<u>Alzheimer’s Disease Diagnostic Guidelines | National Institute on Aging</u> (nih.gov)) for diagnosing Alzheimer’s disease include a standard health evaluation and MMSE evaluations. If indicated, these tests are typically followed with cerebrospinal fluid (CSF) assays for tau and Aβ levels, MRI for brain volume and functionality, and positron emission tomography (PET) scans for Aβ plaques, glucose metabolism, and/or tau fibrils in the brain (Albert et al., 2011; Jack et al., 2011; McKhann et al., 2011; Sperling et al., 2011). These analyses may rule out other dementia etiologies and help to determine disease severity, but they cannot detect AD at its earliest stages or closely predict disease progression, as they do not probe for AD’s earliest biomarkers.

### Current diagnostic methods in development

Spinal taps are invasive, but cerebrospinal fluid assays show promise (Georganopoulou et al., 2005; Toledo et al., 2013b). Nonetheless, assays using CSF analytes have presented challenges with respect to accuracy and reliable disease-state discrimination (Slemmon et al., 2012). More recently, assays for AβO levels in the blood plasma have been developed with promising results (Meng et al., 2019). These assays show a correlation between AβO levels and declining memory scores that appear not to be influenced by age, gender, or ApoE4 status. A promising addition to diagnostic methodology is the detection of AD pathology using targeted *in vivo* brain imaging. The introduction of PET probes for amyloid plaques has been a great technical advance (Klunk et al., 2004) and has established precedent for the usefulness of brain molecular imaging as a diagnostic tool and for proof of efficacy studies in drug development (Johnson et al., 2013). Still, these new imaging tools focus on late-stage by-products of AD such as plaques, rather than early stage instigators such as AβOs.

Prior studies using 5xFAD mice have examined early- and late-stage disease development, but none have looked at the progressive development of AβOs in this model. Here, we present an analysis of memory impairment from 2-9 months of age and the progressive accumulation of AβOs across the same age-span. Our studies presented here use an AβO-selective antibody to characterize the spatiotemporal development of AβOs in the 5xFAD mouse model and demonstrate a correlation with memory impairment. Strikingly, intranasal inoculation of the AβO-selective antibody rescued memory performance in 6-month-old 5xFAD mice. We demonstrate the capability of detecting AβO pathology *in vivo* in the 5xFAD mouse by introducing molecular imaging modalities (MRI and PET) with probes for AβOs. We additionally present immunofluorescent evidence of a remarkable association between AβOs and GFAP-positive reactive astrocytes in the 5xFAD mice. Taken together, we provide further data implicating AβOs as essential diagnostic indicators and therapeutic targets, and show evidence suggesting a mechanism through which AβOs instigate pathological abnormalities: by induction of reactive astrogliosis.

## Materials and Methods

### Materials

ACU193 humanized anti-AβO antibody was a generous gift from Acumen Pharmaceuticals, Inc. Aβ_1-42_ (TFA preparation) was sourced from multiple suppliers (California Peptide, Peptides International, American Peptide). Primary hippocampal cultures were prepared from tissue obtained from BrainBits, LLC, using media and reagents also obtained from BrainBits. All chemicals were purchased from Sigma unless otherwise specified.

### Animals

The 5xFAD Tg mouse model (B6SJL-Tg(APPSwFlLon,PSEN1*M146L*L286V)6799Vas)(Oakley et al., 2006) (Jackson Laboratories) was bred on a non-transgenic background (B6SJLF1/J mice (Jackson Laboratories, RRID: IMSR_JAX:100012)). Aged transgenic and wild-type littermates, 2-20 months old, were used. All mice were kept under a 12/12 h light/dark cycle (7 AM/7 PM) at 22 ± 2 °C. Mice had free access to food and water, including during behavioral experiments, were housed at ≤5/cage (NexGen IVC, Allentown) with enriched environment and daily veterinarian assessment, according to NU’s standard procedures. Procedures complied with NIH’s Guide for the Care and Use of Laboratory Animals (NIH publication No. 80-23, 1996) and were approved by IACUC (protocol #IS00004010). Behavioral experiments were conducted between 12-6 PM.

For intracerebroventricular (icv) experiments, B6SJLF1/J mice (Jackson Laboratories, RRID: IMSR_JAX:100012) were utilized at ages ranging from 6 months of age (30-50 g).

### Aβ Oligomer Preparation

Unlabeled (AβOs) and fluorescently-labeled Aβ oligomers (FAM-AβOs) were prepared essentially according to the protocol published by Klein and colleagues (Lambert et al., 2007; Velasco et al., 2012). Briefly, Aβ_1-42_ (American Peptide or Peptides International) or FAM-Aβ_1-42_ (Anaspec) was dissolved in hexafluoro-2-propanol (HFIP) and distributed into microcentrifuge tubes. Hexafluoro-2-propanol was removed by evaporation and traces removed under vacuum; the tubes were stored at −80°C. For unlabeled AβOs, an aliquot of Aβ_1-42_ was dissolved in anhydrous dimethyl sulfoxide (DMSO) to ~5 mM, and diluted in ice-cold Ham’s F12 medium without phenol red (Caisson Laboratories) to 100 μM. For FAM-AβOs, an aliquot of each peptide was dissolved in anhydrous dimethyl sulfoxide (DMSO) to ~5 mM, mixed 5:1 (mol: mol) Aβ: FAM-Aβ, and diluted in ice-cold Ham’s F12 medium without phenol red (Caisson Laboratories) to 100 μM. For both AβO preparations, this solution was incubated at 4°C for 24 hr. and centrifuged at 14 000 g for 10 min. The supernatant, defined as the AβO or FAM-AβO preparation, was transferred to a clean microfuge tube and stored at 4°C until use. Protein concentration was determined using Coomassie Plus protein assay kit (Pierce).

A modification of this protocol was used to produce crosslinked AβOs (Cline et al., 2019b).

All preparations were tested for quality using SDS-PAGE on a 10-20% Tris-Tricine gel followed by both silver stain and Western blot with NU2 anti-AβO antibody (Lambert et al., 2007; Velasco et al., 2012).

### Cell Culture

Hippocampal cells were prepared and maintained for at least 18 days as previously described (Gong et al., 2003) by using (0.002%) poly-L-lysine coated coverslips plated at a density of 1.04 x 10^4^ cells per cm^2^ in Neurobasal media (Brainbits, LLC) with B27 supplements and L-glutamine (2.5 μM).

### Aβ Oligomer Incubation and Immunolabeling of Cells

Cells were incubated at 37°C in conditioned media collected from the cell cultures containing crosslinked AβOs or FAM-AβOs or an equivalent dilution of vehicle. Following incubation with AβOs or vehicle for 60 min, the cells were rinsed rapidly 3 times with warm media then fixed by adding an equal volume of warm 3.7% formaldehyde (in PBS) to the third rinse in each well/dish and allowing it to sit at RT for 5 min. The media/formaldehyde was completely removed and replaced with a volume of 3.7% formaldehyde for 5 min at RT. Cells were blocked in 10% normal goat serum (NGS) in PBS or HBSS for 45 min at RT then incubated overnight at 4°C on an orbital shaker with fluorescent-tagged antibody or anti-AβO probe diluted in blocking buffer. The cells were washed 3 times for 5 min each with PBS or HBSS. After secondary antibody incubation, coverslips were mounted onto glass slides using ProLong Gold Anti-fade reagent with DAPI (Invitrogen) and imaged using an epifluorescence (TE2000, Nikon), a widefield fluorescence microscope (Leica DM6B, Leica Corp.), or confocal microscope (Leica SP2, Leica Corp).

### AβO intracerebroventricular (icv) administration in mice

Icv injections and behavior testing were performed in 4 independent experiments of 13-21 mice each. Littermates were arbitrarily assigned to different injection groups, targeting 5-10 mice/group for statistical power (n = ((Z_α/2_*)/E)^2^ at α = 0.05; σ = 10.55 and E = 6.67 derived from pilot studies).

Mice were lightly anesthetized (2% isoflurane) during injection (~1 min). AβOs (1, 10 pmol in 3 μl) or vehicles were administered icv free-handed (Bicca et al., 2015). Separate needles were used for each vehicle, progressing from low-high AβO concentration to minimize carryover. No analgesics or anti-inflammatory agents were necessary. Mice were monitored constantly for recovery of consciousness and ambulation, then periodically for food-and-water intake until behavior analysis. Needle placement was confirmed by brain dissection after behavioral experiments (euthanization: CO_2_ then decapitation). Mice showing needle misplacement (3 mice) or cerebral hemorrhage (2 mice) were excluded from analysis; final n = 5-7 mice/group.

### Object Recognition/Location Recognition (NOR/NLR) Tasks

Tasks were performed essentially as described (Bicca et al., 2015), to evaluate mouse ability to discriminate between familiar and new, or displaced, objects within an area, measured by object exploration (sniffing, touching). The open-field testing arena was constructed of gray polyvinyl chloride at 21×21×12” (WxLxH), with a 5×5 square grid on floor and visual cue on wall. 24 h post-injection, mice underwent 6 min sessions of habituation and training, with 3 min between. All sessions were video recorded and analyzed by two researchers blind to experimental groups. During habituation and training, mice were screened for ability to move about the arena and explore the objects, two activities required for accurate memory assessment in subsequent testing sessions. Locomotive inclusion criteria (>100 grid crossings and >15 rearings; evaluated in habituation) were based on extensive previous experiments with the same mouse strain and arena; 3/65 mice did not meet this criterion. During training, mice were placed at the arena center with two objects, which were plastic and varied in shape, color, size and texture. Exploration inclusion criteria were low exploration (<3 sec total) or object preference (>50% of total time for either object); 7 of remaining 62 mice did not meet this criterion.

Hippocampal-related memory function was assessed 24 h post-training by displacing one of the two training objects. Cortical-related memory function was assessed 24 h later by replacing the displaced object with a novel object. Hippocampal-related memory function was re-tested 31-38 days post-injection by displacing the novel object. Memory dysfunction was defined as an exploration of the familiar object for >40% total time. Mice were arbitrarily assessed by cage. The arena and objects were cleaned thoroughly between sessions with 20% (v/v) alcohol to minimize olfactory cues.

### Immunolabeling of slices

Free floating 45 μm thick sagittal sections were cut using a Leica SM2010 R sliding microtome and transferred to sterile TBS for storage. Sections were gathered and placed sequentially into wells (~4 per well). Sections were then randomly selected from each well to perform antibody staining using the primary antibodies ACU193 (0.2 μg/ml), Alexa Fluor^®^ 555-conjugated NU4 (0.92 μg/ml), Cy3-conjugated anti-GFAP (1:800, Sigma) and the secondary antibody Alexa Fluor^®^ 633 goat anti-human IgG (1:2000, Invitrogen). Floating slices were rinsed 3×10 min with TBS and blocked with blocking buffer (10% NGS with 0.3% Triton X-100 in TBS) for 60 min at room temperature. Slices were then incubated with the respective antibodies in blocking buffer overnight at 4°C with gentle rotation. Sections were washed 3 x 10 mins in TBS and incubated with secondary antibody for 3 hours at room temperature (RT) with orbital agitation in the dark. Secondary was prepared in blocking buffer diluted 10-fold with TBS. Sections were then washed 3 x 10 mins in TBS, mounted using ProLong Diamond^®^ antifade mounting media with DAPI (Invitrogen) and 24×60mm No. 1.5 glass coverslips (Thermo Scientific). Z-stacks of the brain sections were collected at 10x or 100x on a Leica SP5 confocal microscope and analyzed with ImageJ.

### Thioflavin S counterstain

Thioflavin-S counterstaining to NU4 immunofluorescence labeling was performed as previously described (Guntern et al., 1992) with a few modifications (Viola et al., 2015). 5xFAD and WT brains were sliced at a thickness of 50μm and immunolabeled following the same protocol described above (immunolabeling of slices). Slices were incubated with antibody as described above. The slices were then washed with PBS for 5 times 5 min each and incubated with 0.002% of Thioflavin-S solution in TBS-T (diluted from a stock solution 0.02% of Thioflavin-S in distillated water) for 10min. Slices were then washed 3 times for 1 min in 50% ethanol and 2 times in TBS-T for 5 min. The slices were mounted with ProLong Gold Antifade reagent for examination by fluorescence microscopy. Images were acquired at 40x magnification and analyzed by ImageJ software.

### Radiolabeling and Quality Control

Antibodies, NU4 and non-specific mouse IgG or ACU193 and non-specific human IgG were radiolabeled with positron emitter ^64^Cu (^64^CuCl2 in 0.1 M HCl; radionuclide purity >99%, Washington University). For radiolabeling, Wipke and Wang’s method was applied (Wipke et al., 2002). Basically, antibodies mentioned above were conjugated with DOTA-NHS-ester (Macrocyclics, Dallas, TX) and then radiolabeled with ^64^Cu.

#### Conjugation

Antibody solutions were buffer exchanged with PBS using YM-30 Centricon^®^ centrifugal filters (Millipore, Billerica, MA). For conjugation, antibodies were reacted with DOTA-NHS-ester in 0.1 M Na_2_HPO_4_ buffer of pH 7.5 at 4°C for 12 - 16 h in a molar ratio of DOTA-NHS-ester:antibody = 100: 1. After conjugation, the reaction mixture was centrifuged repeatedly (5 times) through a YM-30 Centricon^®^ centrifugal filter with 0.1M pH 6.5 ammonium citrate buffer to remove unconjugated small molecules. The concentrations of purified antibody-conjugate was determined by measuring the absorbance at 280 nm in a UV spectrophotometer.

#### Labeling

When labeling with ^64^Cu, 1 mg DOTA-conjugated NU4 and 5 mCi (185 MBq) of ^64^Cu as incubated in 0.1 M ammonium citrate buffer, pH 6.5, at 43°C for 1 hour. Labeled antibody was separated by a size-exclusion column (Bio-Spin6, BIO-RAD Laboratories).

#### Quality Control

Radiochemical purity of antibody was determined by integrating areas on the Fast Protein Liquid Chromatography (FPLC) equipped with a flow scintillation analyzer. This analysis was conducted on a Superpose 12 10/300 GL (Cytiva) size-exclusion column and characterized by the percentage of radioactivity associated with the 150 kDa protein peak. The stability of the ^64^Cu radiolabeled mAbs was determined by bovine serum challenge at 44 hours.

#### Conjugation efficiency

Based on our preliminary data, > 90% of conjugation rate, >70% of labeling rate is achieved by following prescribed protocol.

### Overall details of micro PET and micro CT acquisition

Mice were placed in a 37.5 °C heated cage 20-30 minutes prior to radiotracer injection and moved to a 37.5 °C heated induction chamber 10 minutes prior to injection where they were anesthetized with 2-3% isoflurane in 1000 cc/min O_2_. A dose of 40 μg/200 μCi in 100 μL of proposed PET tracers was administered intravenously through the tail vein. Each animal was administered a dose ranging from 30-40 μg NU4PET, ACU193PET, or non-immune IgGPET. Probes were administered in a single dose. PET/CT imaging was conducted at 0, 4, 24, and 48 h to measure for changes in distribution and time required for probe clearance or decay.

NU4PET scans were acquired using a Genisys^4^ PET (Sofie Biosciences, Culver City, CA) system and CT scans were acquired using a Bioscan NanoSPECT/CT (Washington, D.C.). When scanning, all mice were placed prone on the bed of the scanner. A 10 minute static acquisition was used for PET imaging followed immediately by a 6.5 minute CT acquisition both utilizing the mouse imaging chamber from the Genisys^4^. PET reconstruction was performed without attenuation correction using 3D Maximum Likelihood Expectation Maximization (MLEM) with 60 iterations and CT reconstruction used Filtered Back Projection with a Shepp-Logan Filter. PET and CT reconstructions were exported in dicom image format and fused using custom software developed by the Small Animal Imaging Facility at Van Andel Institute. Fused PET/CT images were analyzed using VivoQuant Image Analysis Suite (inviCRO, LLC, Boston, MA). Standardized Uptake Values (SUV) were calculated using the mouse body weight and corrected for residual dose in the injection syringe and the injection site, as applicable. The formula used to calculate SUV was

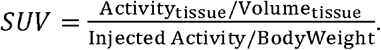

### Evaluation NU4PET (^64^Cu-NU4) in AβOs detection

Two groups (n = 3/ group) of 6 months old 5xFAD Tg AD mouse model and 2 groups (n = 3/ group) WT mouse model were used for evaluating the capability of AβOs detection. NU4PET (^64^Cu-NU4) or non-specific IgGPET (^64^Cu-IgG) was injected into each 5xFAD Tg AD mouse model and WT mouse model groups, respectively.

Target (AβOs)–Background (normal tissue) contrasts in PET images were used to distinguish the difference of the capability of AβOs detection between NU4PET and IgGPET in different mouse models. Tracer uptake of high intensity (hot) areas and background tissues in the brain were chosen by drawing regions-of-interest (ROI) along the edges of the areas from the PET images. Average pixel values of each ROIs were acquired and use in Target (AβOs)–Background (normal tissue) contrasts calculation. The formula used to calculate Target-Background contrast was

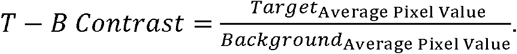

### Tissue Biodistribution Assessment

Animals were sacrificed immediately after the 44 hour post injection image was acquired. Blood was collected, while brains and 13 other organs and tissues were harvested and weighed. After the blood sample was taken from the heart (~500-1000μl), 10 ml of saline was injected into left ventricle while the heart was still beating to flush out the residual blood in the organs. Radioactivity in each tissue (cpm) wa measured using the γ-scintillation counter. Percentages of the injected dose/gram (%ID/g) were calculated for each tissue/ organ by the following formula.

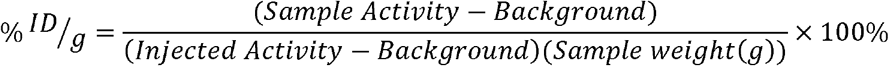

Student’s t-test was conducted to the results between different groups. *P*<0.05 is considered statistically significant.

### Synthesis of Magnetic Nanostructures (MNS)

16 nm magnetite nanoparticles were synthesized by decomposition of iron-oleate at 320°C as described in an earlier report.(Park et al., 2004)

#### Synthesis of Iron-oleate complexes

10.8 g of iron (III) chloride hexahydrate and 36.5 g sodium oleate were dissolved in a mixture of 60 ml distilled water, 80 ml ethanol and 140 ml hexane and heated at 60°C for 4 hr. The organic layer of the biphasic mixture becomes dark, indicating phase transfer of iron (III) ions and formation of iron oleate complex. The resulting dark solution is separated and washed with water three times.

#### Synthesis of 16 nm magnetite nanoparticles

18 g of iron oleate complex and 2.58 g of oleic acid were dissolved in 100 g of octadecene at room temperature and heated to 320°C at a rate of 3.3°C per minute. The reaction mixture is kept at 320°C for 40 min., then cooled down to room temperature. Resulting nanoparticles are separated from the solution by addition of ethanol and ethyl acetate followed by centrifugation.

##### Preparation of Dopamine-TEG-COOH and Phase Transfer

To make the organic phase synthesized MNS suitable for biological application, we functionalized the MNS using an in-house synthesized ligand with carboxylate as terminal group (for antibody conjugation), tetraehylene glycol(TEG) as a stabilizer, and nitrodopamine (nDOPA) as an anchor due to its high affinity for Fe (Nandwana et al., 2016).

Synthesis of carboxylate terminated nDOPA ligand and functionalization of the MNS was carried out according to the following protocol. Tetraethylene diacide, N-hydroxysuccinimide (NHS), N,N’-Dicyclohexylcarbodiimide (DCC), nDOPA hydrochloride and anhydrous sodium bicarbonate was dissolved in chloroform under argon atmosphere and stirred for 4 hr. Hexane stabilized MNS were added and stirred for another 24 hr. The precipitate formed was separated by magnet, dispersed in water and purified by dialysis.

##### Conjugation of antibody to MNS

The conjugation of buffer stabilized MNS with antibody was done using a conventional carboxyl-amine crosslinking method. We first activated the carboxyl terminated MNS by sulfo-N-hydroxy succinimide (SNHS) and 1-Ethyl-3-(3-dimethylaminopropyl)carbodiimide (EDC) followed by incubation with corresponding antibody (NU4 or IgG1, with or without fluorescent label) overnight. Conjugated MNS were separated by magnet to remove excess reagent and antibody then re-dispersed in working media. Conjugation efficiency was estimated using UV spectroscopy (absorbance at 280nm) of the magnetically separated supernatant.

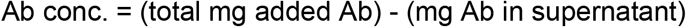

### Intranasal immunization

Mice were anesthetized with isoflurane and then placed on their backs with their heads positioned to maximize the residency time for the delivered material to remain on the olfactory surface. Each naris was administered with ACUMNS or non-immune IgGMNS (10 μl/naris), using a sterile micropipette, slowly over a period of 1 min, keeping the opposite naris and mouth closed to allow complete aspiration of delivered material. Steps were repeated up to 5 times, maintaining anesthetization in between inoculations, for maximum doses of up to 50μl/naris

### Magnetic Resonance Imaging of Tg and WT mice in vivo

Following intranasal inoculation, the probe was allowed to distribute for 4 hours before MR imaging was performed according to imaging methodology described in Mundt et al.(Mundt et al., 2009) T1, T2, and T2* weighted MR images were acquired on a Bruker BioSpec 9.4T magnet, using a 25 mm RF quadrature coil. The in-plane resolution was 75 μm with slice thickness 0.4 mm. T1- and T2-weighted images provide anatomical guidance as well as some localization of the ACUMNS and were acquired with a fat suppressed spin echo sequence (Rapid Acquisition with Relaxation Enhancement, RARE) with the following parameters for T1-weighted (TR=1000 ms, TEeff=13.2 ms, rare factor 2, number of excitations, NEX=4) and for T2-weighted (TR=3500 ms, TEeff=58.5 ms, rare factor 4, NEX=4). T2*-weighted imaging provides more of the localization of the NU4MNS as the iron causes local changes in magnetic susceptibility which T2* weighted images can be sensitive to. A gradient echo sequence was used with the following parameters (gradient echo fast imaging, GEFI; TR=1200 ms, TE=5.6 ms, flip angle 35° and NEX=4).

## Results

### Memory dysfunction in 5xFAD mice begins shortly after AβO emergence and progressively worsens with concomitant AβO accumulation in the hippocampus

#### Tg 5xFAD NOR/NLR

Amyloid plaque development and intraneuronal Aβ42 accumulation are well-established in the 5xFAD transgenic (Tg) mouse model of Alzheimer’s disease. There is robust plaque buildup around 5-6 months of age (Ohno et al., 2006) and intraneuronal Aβ42 accumulation begins as early as 2 months (Oakley et al., 2006). The majority of neuropathological studies in 5xFAD mice have used probes that show amyloid plaque development; how 5xFAD memory impairment coincides with AβOs pathology and development is much less well-characterized. In order to characterize how memory loss correlates with AβOs in the 5xFAD mice, we used the well-established novel object recognition task (NOR) for non-spatial (cortical) memory (Cohen and Stackman, 2015; Denninger et al., 2018) and the novel location recognition task (NLR) for spatial (hippocampal) memory (Antunes and Biala, 2012; Bengoetxea et al., 2015; Grayson et al., 2015; Denninger et al., 2018). We assessed memory in mice aged 2-18 months. 5xFAD mice showed no evident memory impairment at 2 to 3 months old (Figure 1a). By 4 to 5 months old, most transgenic mice showed memory impairment, and by 6 to 7 months of age memory impairment was apparent in all 5xFAD mice. Importantly, at 4 months old, the majority of 5xFAD mice were impaired in both the hippocampal-dependent and cortical-dependent tasks; there were, however, some mice that showed only cortical-impairment. Though less obvious than their Tg littermates, memory loss was detected at 9 months of age in wild-type mice. In summary, we showed that 5xFAD mice first present memory impairment between 3 and 4 months of age. This memory dysfunction afflicts more mice as their age increases until, at 6 to 7 months, all of the Tg mice are impaired in both hippocampal-dependent and cortical-dependent tasks. These data indicate that memory impairment begins before observed amyloid plaque build-up in the 5xFAD mice.

**Figure 1.**
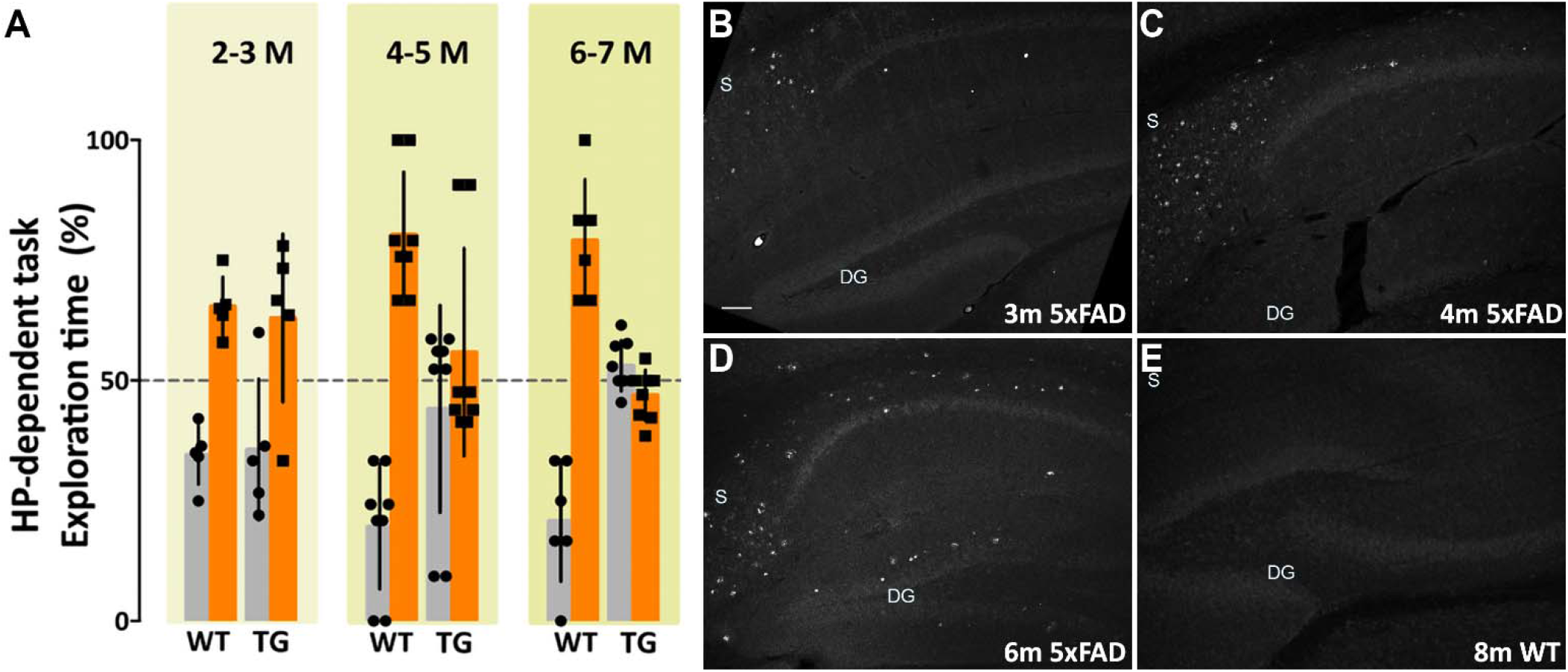
Memory dysfunction in 5xFAD mice is substantial by 4 months and is preceded by AβO pathology, detectable by 3 months of age. (Left) 5xFAD mice and wild-type littermates were assessed for memory dysfunction using novel location recognition (NLR; hippocampal-dependent task) and novel object recognition tasks (NOR; cortical-dependent task). Ages ranged 2-12 months. Data shown here are for the hippocampal-dependent NLR assay. In 5xFAD mice, memory impairment was negligible at 2-3 months, substantial by 4-5 months, and fully penetrant by 6 months of age. Statistical analysis shows that there was no significant difference between the behaviors of the WT mice and the 5xFAD mice at ages 2-3 months, but a statistically significant difference was evident between the recognition task behaviors of the WT mice and 5xFAD mice for ages 4-5 months (p<0.001) and 6-7 months (p<0.0001). (Right) Sagittal brain sections were obtained from 5xFAD and WT mice at ages 2, 3, 4, 6, and 8 months and probed for AβO pathology using a humanized AβO monoclonal antibody. Fluorescent signal was barely detectable at 2 months of age in some mice, more readily detectable by 3 months in all Tg mice, and robust by 6 months. Wild-type littermates presented no signal. Scale bar = 100 μm.

#### Immunohistofluorescence validation of AβO development

The development of amyloid plaque pathology is well-established in the 5xFAD mouse model (Oakley et al., 2006; Ohno et al., 2006). Amyloid plaques, however, are no longer considered the most germane Aβ species to AD pathology (Overk and Masliah, 2014; Viola and Klein, 2015; Selkoe and Hardy, 2016; Cline et al., 2018; Li and Selkoe, 2020). Characterizing the development of the most relevant species, putatively AβOs, and their association with other pathological changes in AD, such as glial activation or pTau accumulation, is necessary to better understand disease progression in this model. Sagittal sections of brain tissue, collected and fixed from WT and 5xFAD mice at ages 2, 3, 4, 6, and 8 months of age, were immunolabeled with ACU193 and imaged using confocal microscopy. ACU193, a humanized monoclonal antibody that targets AβOs, has been shown to selectively bind oligomers *in vitro* (Krafft et al., 2013; Goure et al., 2014; Savage et al., 2014) and in the TG2576 mouse model. Here, using ACU193 to probe for AβOs, we show the progressive, spatio-temporal accumulation of AβOs in the hippocampus of 5xFAD mice (Figure 1b). AβOs first appear in the subiculum as early as 2 months of age in some mice and are detectable by 3 months in all 5xFAD Tg mice examined. In the transgenic mice, AβOs show a continued accumulation in the subiculum and a spreading of pathology to CA1, CA2 and the dentate gyrus. This timing suggests that AβOs are associated with the observed memory loss.

#### ACU193 detects AβOs bound to primary neurons with high specificity

To validate the specificity of ACU193 for AβOs, the antibody was used *in vitro* to detect synthetic preparations of oligomers introduced into primary hippocampal neurons in culture (Supplemental Figure 1). Primary hippocampal neurons were treated with cross-linked AβOs, which have been shown to preserve AβO structure *in vitro* (Cline et al., 2019b), or vehicle control. The cells were subsequently fixed and labeled with ACU193 at increasing dosages. Confocal imaging of the cells showed somatic staining of AβOs in addition to small, nanoscale puncta along dendritic processes (labeled with MAP2). These ACU193-positive puncta are likely AβOs binding to dendritic spines, as seen in previously published work (Lacor et al., 2007; De Felice et al., 2009; Pitt et al., 2017). Minimal ACU193 labeling was observed on vehicle-treated neurons, indicating its specificity for AβOs.

### ACU193 and NU4 detect AβOs

Additional support for the specificity of ACU193 can be seen in comparing the distribution of ACU193 in brain sections with the distribution of NU4, a well-established AβO monoclonal antibody (Lambert et al., 2007; Xiao et al., 2013; Viola et al., 2015). Using ACU193 and NU4 conjugated to Alex Fluor 555 we found that both antibodies similarly detected AβOs in the subiculum and other areas of the hippocampus (Figure 2) including CA1, CA2 and the dentate gyrus. ACU193-(cyan) and NU4-positive (magenta) cells were observed accumulating in a nearly identical pattern, from 3 months to nine months of age. ACU193 and NU4 selectively detect AβOs in the 5xFAD mice with virtually no signal in WT mice.

**Figure 2.**
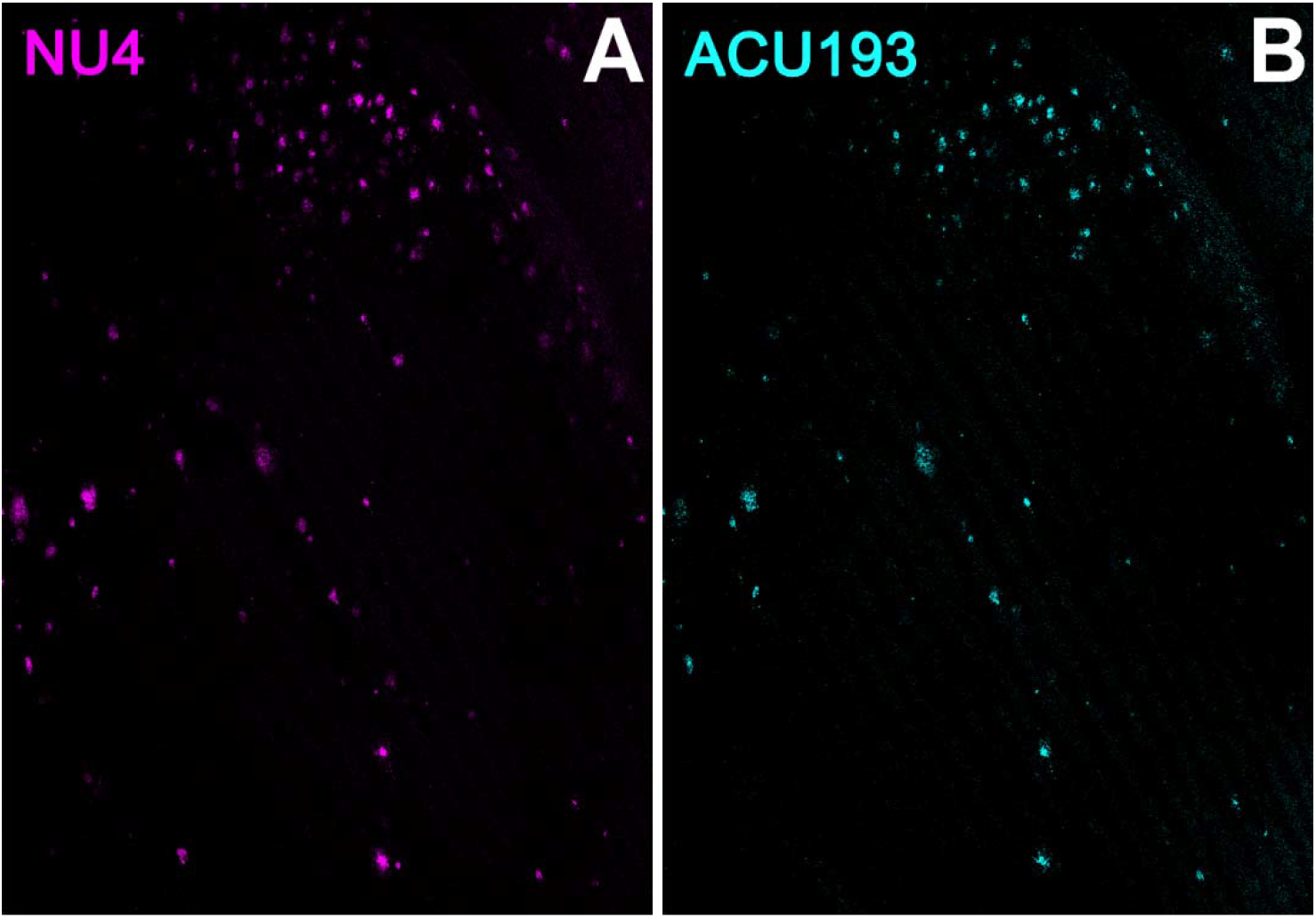
ACU193 and NU4 detect AβOs *ex vivo*. Sagittal sections from 9-month-old 5xFAD mice were immunolabeled with 2 different anti-AβO antibodies, NU4 and ACU193, to determine the extent to which AβO pathology is detected by both antibodies. Data show that AβOs accumulate and that ACU193 and NU4 show very similar detection of AβOs.

### Alzheimer’s-associated astrocyte pathology develops concomitantly with AβOs

To determine whether other Alzheimer’s related pathologies show developmental regulation or accumulation in the 5xFAD mouse model for AD in association with AβOs, we examined immunohistochemical patterns of glial fibrillary acidic protein (GFAP), activated microglia (Iba1), and phosphorylated tau (pTau). Immunolabeling for pTau yielded difficult to interpret results which varied amongst the different antibodies for the same epitope and often did not match the literature. Instead, we focused on the inflammatory pathways, stimulated by the strong interest in the involvement of inflammatory responses in AD, in particular a new and growing interest in astrocytes (Wang et al., 2021). Immunolabeling for activated microglia (Iba1) (Supplemental Figure 2) indicated that the WT mice have more ramified microglial cells (resting) while 5xFAD littermates have more amoeboid and activated-shaped microglial cells. Notably, microglial activation was evident at 2 months, with no obvious increase in abundance seen in older animals. In contrast, sagittal sections from 5xFAD or wt mice, aged 3-9 months, were immunolabeled with antibodies against GFAP and co-labeled with ACU193, then imaged by confocal microscopy. We found a marked spatiotemporal association of GFAP pathology with ACU193-positive AβOs in the 5xFAD mice. GFAP (Figure 3, magenta) pathology first appeared in the subiculum at 3 months of age concurrent with the first appearance of AβOs (cyan) in the subiculum and in close proximity to one another. As the mice aged, GFAP and ACU193-positive pathology concomitantly spread throughout the subiculum and hippocampus (Figure 3, B & E). At 9 months, WT mice have minimal GFAP expression (Figure 3C) and no AβOs (Figure 3F). These patterns are consistent with possible induction of reactive astrogliosis by AβOs. At higher magnification, we observed GFAP-positive reactive astrocytes surrounding an ACU193-positive neuron and projecting their processes onto the cell soma (Figure 3I). In addition, we observed micron-wide ACU193-positive puncta adjacent to astrocytic processes distant from the cell soma.

**Figure 3.**
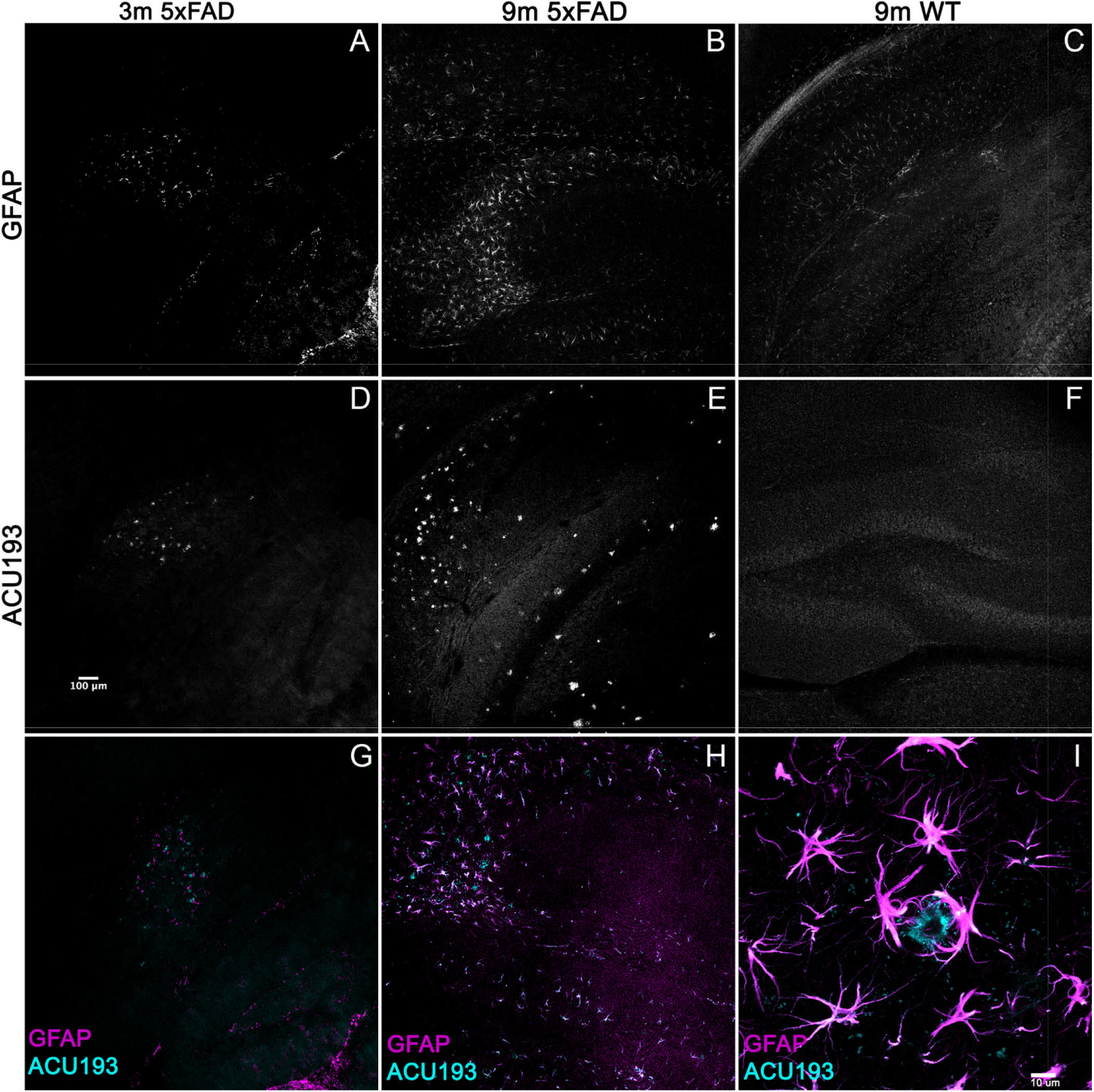
Alzheimer’s-associated astrocyte pathology develops concomitantly with AβOs. Sagittal sections from 5xFAD mice, aged 3-9 months, and their wild-type littermates were immunolabeled with antibodies against GFAP and ACU193, then imaged on the Leica SP5 confocal microscope at 10x and 100x. Data show that, like the ACU193, GFAP positive glial cells accumulate in an age dependent manner. Sale bar = 100 μm for panels A-H ad 10 μm for panel I.

### AβOs given to WT littermates induces memory impairment within 24 hours

#### ICV AβOs induce impairment in NLR/NOR

While the previous data indicate a relationship between AβO accumulation and memory dysfunction in the 5xFAD mice, the question remained whether AβOs cause the observed memory loss. We therefore asked whether injection of AβOs into WT littermate mice would induce similar behavioral dysfunction. Wild-type littermates from the 5xFAD colony were injected with either 10 pmol synthetic AβOs or volume equivalent of vehicle control into the right lateral ventricle, following our previously established protocol (Lambert et al., 2007; Velasco et al., 2012; Cline et al., 2019b). After 24 hours, the mice were assessed by the NLR task, and later, the NOR assay at 48 hours post-injection. We found that ICV injection of AβOs induce memory dysfunction within 24 hours and impacts both cortical (NOR) and hippocampal (NLR) memory (Figure 4). As in the 5xFAD mice, AβO injected mice showed no preference to either new or old objects and explored both equally. Vehicle-injected mice scored no different from wild-type in these tasks. These data show that AβOs are sufficient to induce memory impairment within 24 hours post-injection in wild-type mice. We next sought to establish the functional effect of neutralizing these AβOs in the 5xFAD mice.

**Figure 4.**
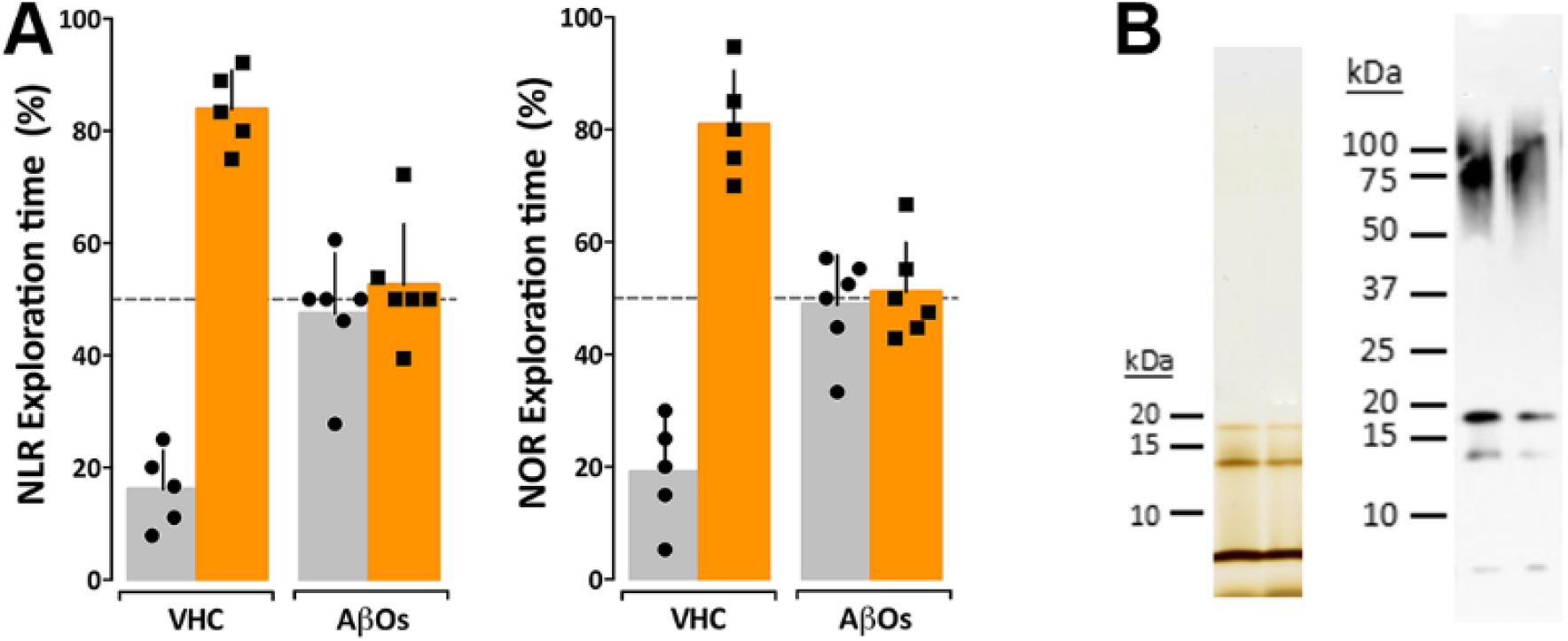
Intraventricular AβO injection causes memory impairment in wild type mice within 24 hours. (A) Wild type mice were tested for performance in recognition tasks beginning 24 hours after receiving vehicle (VHC) or AβO injections (AβOs) (10 pmols in 3 μl) into the right lateral ventricle. Mice first were assessed for novel location recognition (NLR; 24 hr post-injection) and subsequently for novel object recognition (NOR; 48 hr post-injection). AβO-injected mice were unable to perform either recognition task. Statistical analysis shows that there is a statistically significant difference between the recognition task behaviors of the WT mice and the AβO injected mice (p<0.0001). (B) Silver stain (left) and Western blot (right) analysis of the AβOs used for injections and other assays in this study shows preparations contain trimer, tetramer, and higher molecular weight species as has been shown before (Lacor et al., 2007; Lambert et al., 2007; Velasco et al., 2012).

### Oligomer-selective antibodies engage and neutralize AβOs responsible for memory dysfunction in 5xFAD mice

#### ACU193-based probes ameliorate memory dysfunction

We have previously observed no short-term detrimental impact after inoculation of our AβO antibodies into 5xFAD mice, but no studies have been done to determine the long-term positive or negative effects in these mice. To determine the impact of AβO-neutralization in 5xFAD mice, 6- and 7-month-old mice were first assessed for memory impairment using the NLR/NOR assay. Mice were then inoculated with ACU193-based probes and imaged 24 hours later *in vivo* to ensure target engagement (see next section). The mice were then housed for 30-40 days to monitor any adverse effects or changes in behavior before being reassessed for memory impairment in the NLR/NOR tasks. Strikingly, we found that 6-month-old 5xFAD mice inoculated with the ACU193-based MRI probe had reversal of memory dysfunction, with performance the same as WT controls in the NOR task 30 days post-inoculation (Figure 5). The ACUPET probe similarly ameliorated memory impairment, measured 40 days post-injection. As controls, 5xFAD mice injected with human IgGMNS or IgGPET probe showed no memory improvement. Results from 4 trials of 10-12 animals each show that the ACU193 antibody engages AβOs *in vivo*, completely reversing memory dysfunction in the 5xFAD mice with no evidence of health issues or side effects. The data establish AβOs as the primary instigators of cognitive dysfunction in 5xFAD mice and support the therapeutic relevance of AβO-selective probes.

**Figure 5.**
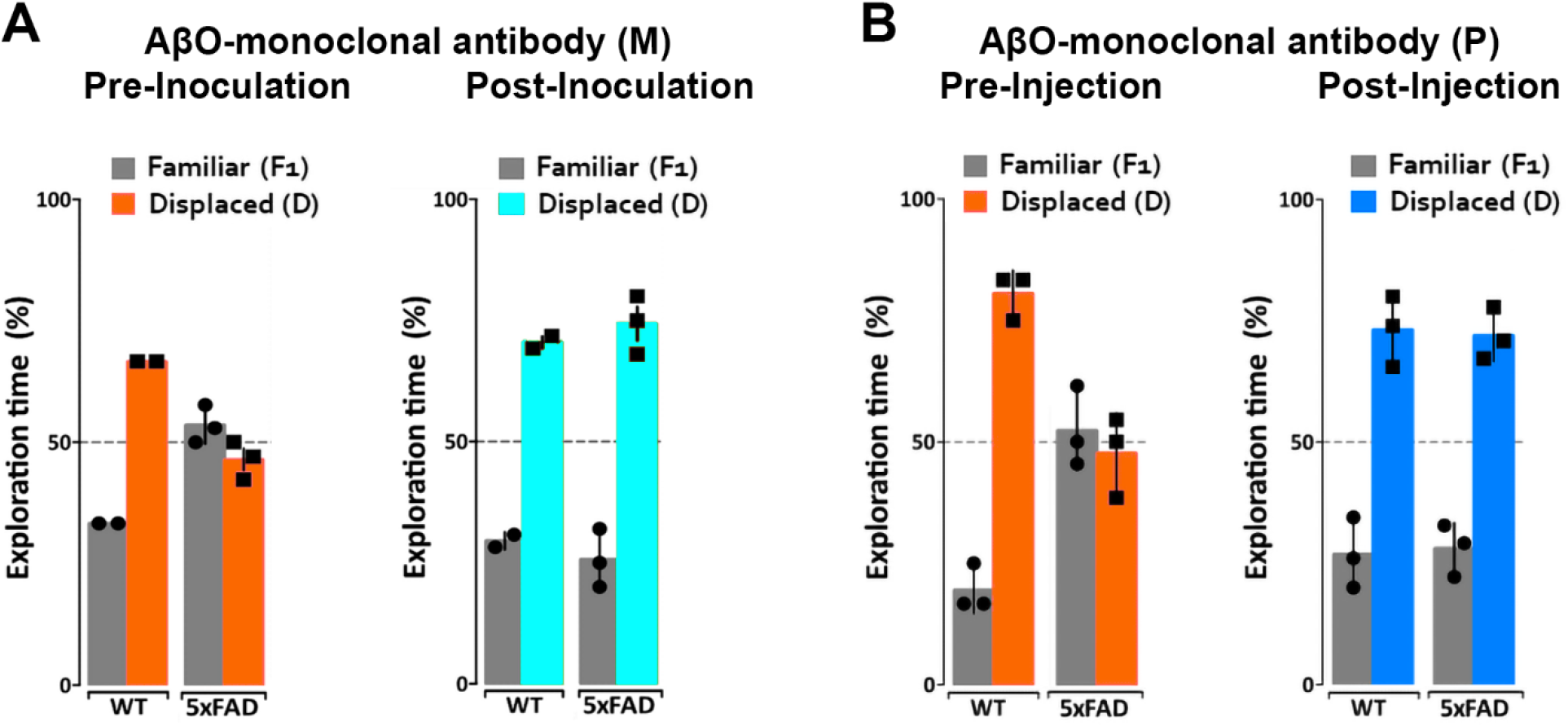
ACUMNS delivered intranasally or ACUPET given iv each rescue memory function in 6- to 7-month-old mice. Tg and WT mice, aged 6 months (A), were tested by NLR and NOR assays to ensure predicted behavioral deficits. Mice were then intranasally inoculated with ACUMNS and imaged for probe distribution and detection of AβO pathology *in vivo*. After imaging, animals were monitored for 30 days for signs of adverse reactions to the probe (none detected), then re-tested by NOR. The 6-month-old animals showed a significant recovery of memory impairment 30 days after inoculation. Human IgGMNS showed no impact on memory recovery. (B) To test the impact of the ACUPET probe on memory function, Tg and WT mice, aged 7 months, were tested by NLR and NOR assays prior to imaging as before. Mice were then injected, via tail vein, with ACUPET or non-specific IgGPET and imaged for up to 24 hours to monitor probe distribution. After imaging, animals were monitored for 40 days for signs of adverse reactions to the probe. Animals were re-tested by NOR at 40 days recovery. 5xFAD animals injected with ACUPET showed a persistent recovery of memory impairment that was not seen in the 5xFAD animals injected with IgGPET. ACU-based probes have no impact on wt behavior. Results are representative of 4 separate trials that showed beneficial impact of these antibody-based probes on memory.

### AβOs imaged *in vivo* using ACU193-based probes distinguish 5xFAD from wild-type mice

#### MRI signal from ACUMNS distinguishes 5xFAD from wild-type mice

Our previous work showed that AβOs can be detected *in vivo* in the 5xFAD mouse model using antibody-based MRI probes which were conjugated to magnetic nanostructures (MNS) (Viola et al., 2015). These prior studies used NU4 as the AβO-targeting antibody, which as shown above, binds similarly to ACU193. Here we show that ACU193 can also be developed into a molecular probe for AβO detection *in vivo*. After baseline imaging by MRI, 12-month-old mice were intranasally inoculated with MNS-conjugated ACU193 and allowed to recover overnight (about 16 hours) before imaging again (Figure 6). MRI data shows an accumulation of the ACUMNS probe in the hippocampus and cortex of the 5xFAD mice that is absent in WT controls. ImageJ quantification of signal intensity in the hippocampi of inoculated mice shows a ~ 30-fold increase in 5xFAD mice over their WT littermates. Using the ACUMNS probe in 18-month-old mice showed similarly robust AD-dependent MRI signal in the hippocampus of the 5xFAD animals, but signals obtained in younger animals (6-months old) were less consistent. These data add to previous studies with the NU4 probe and show that non-invasive *in vivo* imaging of AβOs is possible using the ACUMNS probe, suggesting its potential diagnostic value and ability to confirm target engagement.

**Figure 6.**
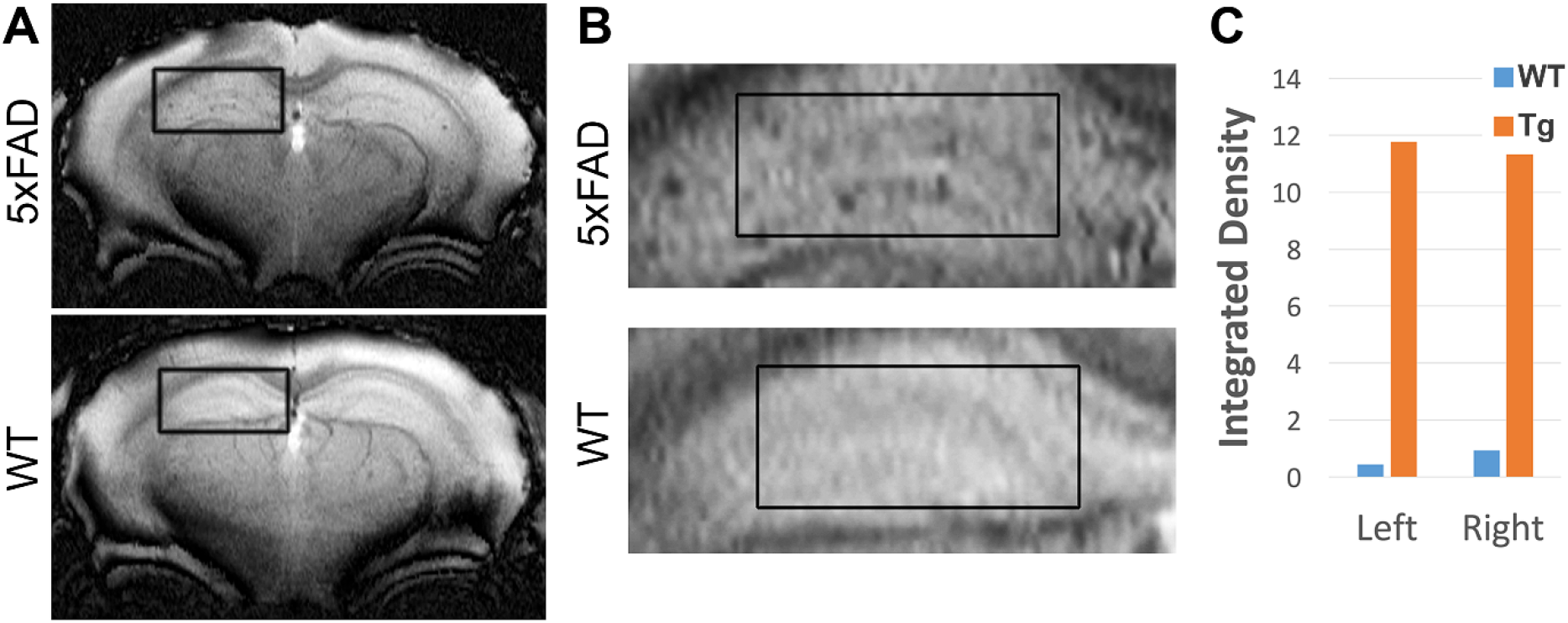
ACUMNS gives AD-dependent MRI signal in hippocampus of 12-month-old 5xFAD mice. *In vivo* studies with ACUMNS probe show robust AD-dependent MRI signal in the hippocampus of 12 month-old mice.

#### Development of an ACU193-based PET imaging probe for early AβO detection

While the spatial resolution of MRI is excellent, its sensitivity is lower than other imaging modalities such as positron emission tomography (PET). Given PET sensitivity is at least 100 times greater than MRI, we thought it might detect very low levels of AβOs during early stages of AD development. ACU193 was conjugated to DOTA, a chelator, as the initial step in the PET probe development. To ensure that this conjugation did not interfere with the antibody’s ability to target AβOs, sagittal brain slices from 5xFAD mice were probed with the ACU193-DOTA probe and counterstained with Thioflavin S (ThioS) for amyloid plaques (Supplemental Figure 3). Results show that ACU193-DOTA detected AβOs in the 5xFAD brain and did not co-localize with ThioS, consistent with previously obtained results showing that ACU193 does not bind amyloid plaques cores (Cline et al., 2019a).

#### ACUPET detects pathology in the brains of 4-month and older 5xFAD mice

The next step was to determine if radiolabeled ACU193-DOTA (ACUPET) detects AD-related AβOs in the 5xFAD mouse brain at an early age. ACU193-DOTA was incubated with ^64^Cu and free isotopes were removed prior to tail vein injection into mice of either 4 or 18 months old, Mice were then imaged at 1, 4, and 24 hours post-injection for ACUPET distribution. At 4 hours post-injection, ACUPET accumulation in the brain was detectable, but not robust. By 24 hours, accumulation of the ACUPET probe in the brains of the 5xFAD animals was evident in both the 4-month-old animals (Supplemental Figure 4A) and the 18 month old animals (Supplemental Figure 4B-D). Animals at 6, 7, 8 and 12 months were also examined and similarly were able to distinguish 5xFAD from WT mice (data not shown).

### AβOs are specifically detected in vivo by NU4PET

#### NU4-based PET probe development

Given the success of the NU4-based MRI probe (Viola et al., 2015), an NU4-based probe was synthesized for PET imaging. NU4 was conjugated to DOTA and tested to ensure that this conjugation did not interfere with the antibody’s ability to target AβOs. Primary hippocampal neurons, pre-treated with fluorescently conjugated AβOs (FAM-AβOs) and were probed with NU4-DOTA (Supplemental Figure 5). Data show that nearly all FAM-AβOs (magenta) were also labeled with the NU4-DOTA probe (colocalization seen as dark blue) and no free NU4-DOTA (cyan) was detected. Vehicle treated cells showed no NU4-DOTA binding. Data confirm the specificity of the NU4-DOTA probe for AβOs, necessary for its use for *in vivo* imaging.

#### NU4PET detects AD-related pathology *in vivo* in 5xFAD mice, distinguishing them from WT

Validation of the AβO-PET probes as effective for early AD diagnostics requires verification that they produce an *in vivo* signal that depends on the presence of AβOs. To validate our new probe, NU4 (Lambert et al., 2007; Acton et al., 2010) and non-specific IgG antibodies were conjugated to DOTA and then radiolabeled with positron emitter ^64^Cu using Wipke and Wang’s method (Wipke et al., 2002). Our next step was to image for AβOs by PET following probe delivery. Animals (12 total), 7 months of age, were injected via tail vein with either NU4PET or IgGPET and then imaged at T=1, 2, 4, 8, 20, 30, 40, and 44 hours after injection. After 44 hours, the animals were euthanized and their brains removed for a final *ex vivo* image of all 12 brains simultaneously (3 animals per group). Results showed the NU4PET specifically identified 5xFAD animals (Figure 7). No signal was detected in all three control groups (5xFAD with IgGPET;WT with NU4PET; WT with IgGPET).

**Figure 7.**
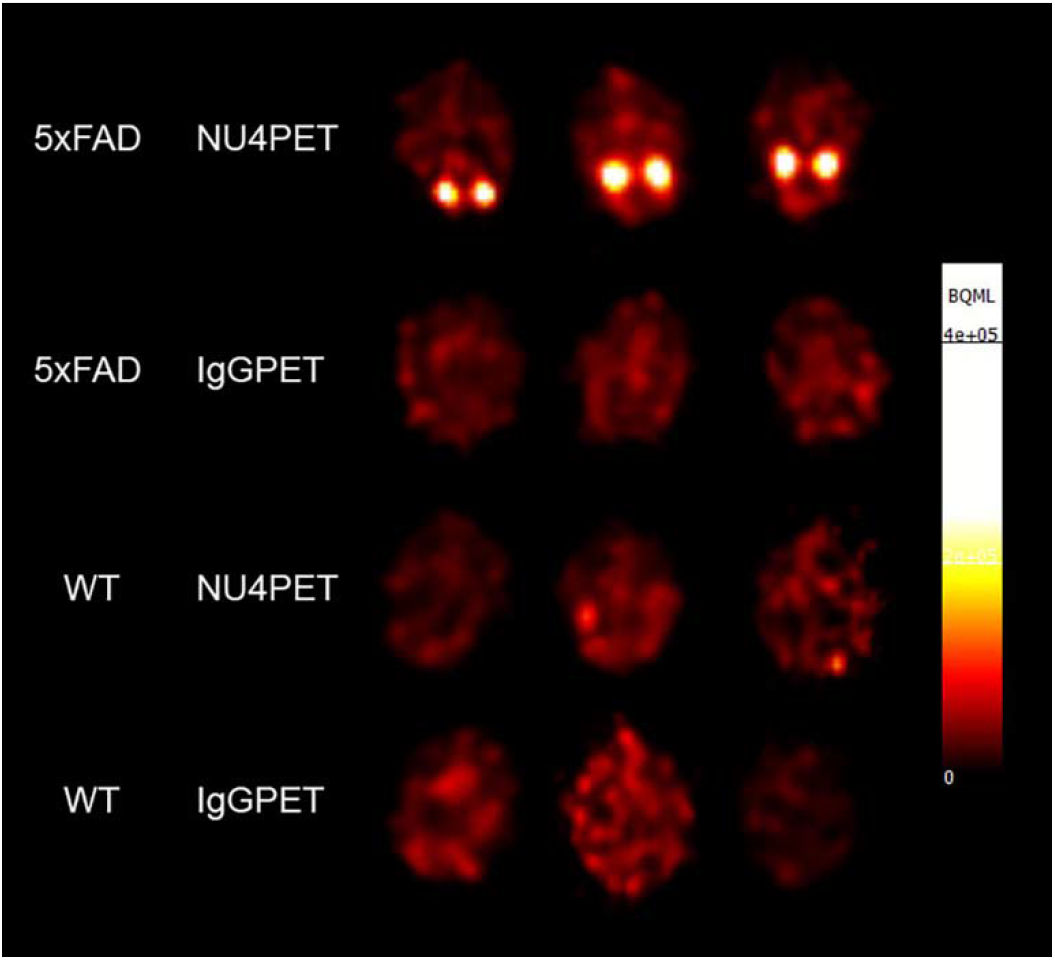
NU4PET probe gives 5xFAD-specific CNS signal. Signal obtained after IV injection of NU4PET showed probe accumulation in the hippocampus of 5xFAD mice (aged 5-7 months). Controls (IgGPET in AD mice; NU4PET in wild type littermates; IgGPET in wild type littermates) showed no signal (3 animals per group).

The fraction of NU4PET probe retained (Supplemental Figure 6) showed good uptake into the brains of the 5xFAD mice but not the WT littermates (quantification of uptake; see Methods). For all mice, the IgGPET probe showed negligible signal. Quantification showed uptake into the brain was comparable to levels of uptake seen with the commercially available Pittsburgh Compound B (PiB) tracer (Mathis et al., 2003; Klunk et al., 2004). To corroborate the presence of AβOs in the animals used for these studies, we analyzed the brain tissue with immunofluorescence. After final PET imaging, the brains were fixed and stored in 10% sucrose until no longer radioactive. Brains were then sliced sagittally at 50 μm and probed with ACU193. Images were collected and analyzed for ACU193 signal intensity (Supplemental Figure 7). Data showed that only 5xFAD mice, and not WT littermates, had AβO pathology. Results confirm the NU4 PET probe gives a signal selective for AβO-positive mice.

## DISCUSSION

Alzheimer’s disease is costly and marked by accumulation of pathological hallmarks such as amyloid plaques and neuronal tangles of hyperphosphorylated tau. Because Aβ plaques have shown poor correlation with AD progression, there has been a rise in the exploration and development of therapeutics that are not based on amyloid (Cummings et al., 2021). This shift in focus has resulted in numerous potential therapies that have made it into clinical trials, but so far there have been limitations on their impact. As an alternative, focusing on AβOs as the target for diagnostics and therapeutics appears to be a promising strategy for developing disease modifying treatments and early diagnosis. Here, we confirm that AβOs can induce memory dysfunction in wild type mice and that AβOs build up in 5xFAD mice in a manner concomitant with astrocyte pathology and with memory dysfunction. Importantly, targeting this buildup with AβO-selective antibodies rescues memory performance. Furthermore, we demonstrate that antibody-based brain imaging probes that target AβOs can be used to identify animals that present with AD pathology, indicating the value of AβO-selective antibodies both for diagnostics and therapeutics.

Recent interest in inflammatory processes and their involvement in AD has grown. Our data showed a striking association between GFAP-positive astrocytes and ACU193-positive AβOs. This association and concomitant increase indicates a potential mechanism for AβO-induced behavioral abnormalities. These findings are particularly intriguing given recent studies indicating AD’s dependence on astrocytes (Huang et al., 2017; Monterey et al., 2021; Nisa et al., 2021; Preeti et al., 2021; Zhou et al., 2021). One especially interesting study showed that when apolipoprotein E (ApoE), a protein expressed in astrocytes which AβOs associate with at synapses, was knocked out in astrocyte-only populations of P301S mice, AD pathology markedly improved (Wang et al., 2021). As ApoE4 is the greatest genetic risk factor of late onset AD, we propose that it may mediate AβO-induced reactive astrogliosis and the subsequent neuropathology instigated by reactive astrocytes. Another study showed that astrocytes were activated into their reactive state via the JAK/STAT3 pathway in 6 month-old 5xFAD mice (Choi et al., 2020). Consistent with the idea that reactive astrogliosis is necessary for behavioral dysfunction in 5xFAD mice, STAT3 phosphorylation inhibition restored cognitive function in the 5xFAD mice. Taken together with our data, we propose that AβOs may induce JAK/STAT3 pathway-dependent reactive astrogliosis in astrocytes which is necessary for observed cognitive dysfunction in 5xFAD mice. In addition to astrocytes, microglia play a major role in AD pathology. The Triggering Receptor Expressed on Myeloid cells 2(TREM2)-expressed in microglia - has already been shown to be involved in AD, with mutations being neuroprotective and TREM2 accumulation being detected in AD patients (Jiang et al., 2013; Benitez et al., 2014; Guven et al., 2020). Previous studies have shown that AβOs associate with TREM2 (Zhao et al., 2018; Zhong et al., 2019; Price et al., 2020), but TREM2 has no impact on established pathology (Yuan et al., 2021).

While interest increases in alternatives to the Amyloid Hypothesis, we are still left with no effective diagnostic tools for identifying AD at its earliest stages when therapeutics have the greatest impact. Currently recommended tests may rule out other dementia etiologies and help to determine disease severity, but they cannot detect AD at its earliest stages or closely predict disease progression. While AD diagnosis has significantly improved with the incorporation of a multiple assay evaluation currently being recommended, the tests still cannot predict disease progression or diagnose AD at its earliest stages because they are not quantifying the earliest biomarkers of the disease. However, alternative detection assays are being developed. Pre-tangle Tau, thought to be the toxic form of tau, has now been detected in MCI and AD and has been found to be one of the earliest tau lesions that correlates with cognitive status (Mufson et al., 2014). Synapse loss (Bastin et al., 2020; Buchanan et al., 2020; Camporesi et al., 2020; Mecca et al., 2020; Pereira et al., 2021), changes in hormone levels (Cheng et al., 2021), changes in blood biomarker levels (Guzman-Martinez et al., 2019; Montoliu-Gaya et al., 2021), electroencephalogram (EEG) readings (Hulbert and Adeli, 2013; Siwek et al., 2015; Lin et al., 2021), retinal assays (Ashok et al., 2020; Mirzaei et al., 2020), and changes in specific protein levels (Buchanan et al., 2020; Colom-Cadena et al., 2020) are some of the myriad assays being developed to try to detect AD earlier and predict when and if the change from mild cognitive impairment (MCI) to AD will occur (Zhang et al., 2021b). All of these new developments are focused towards enabling earlier therapeutic intervention when chances for success would be greatest.

AβOs as a diagnostic resource are currently unavailable. Cerebrospinal fluid assays show promise (Georganopoulou et al., 2005; Toledo et al., 2013a; Savage et al., 2014; Yang et al., 2015; Yang et al., 2019), but spinal taps are invasive and assays using CSF analytes have presented challenges with respect to accuracy and reliable disease-state discrimination (Slemmon et al., 2012). Other assays for AβO levels are under development and show promise as well (Meng et al., 2019). For example, AβO quantification in blood plasma shows a correlation between AβO levels and declining memory scores that appear to not be influenced by age, gender, or ApoE4 status. Recently, the examination of soluble cortical extracts by ELISA found a link between the ratio of AβOs and fibrils with disease. “The ratio of AβO levels to plaque density fully distinguished demented from non-demented patients, with no overlap between groups in this derived variable.” (Esparza et al., 2013)

Because AβOs are regarded as the first toxin to appear in disease progression, they should provide an excellent target for diagnostic imaging (Hefti et al., 2013; Goure et al., 2014). The usefulness of targeting AβOs is indicated by human neuropathology studies in which AβOs initially appear bound to discrete neurons, localizing to synapses in dendritic arbours (Lacor et al., 2004) through putative association with clustered cell surface receptors (Ferreira and Klein, 2011). FAM-AβOs bind at discrete sites on dendrites, showing saturable, concentration-dependent synaptic binding (Viola et al., 2015), further suggesting their potential as a suitable target for an antibody-based diagnostic probe. Pronucleon™ imaging used engineered peptides that deliver a readout when associated with beta-rich Aβ fibers and oligomeric Aβ (Nyborg et al., 2013). Several PET probes have also been developed including a probe from curcumin^18^F (Rokka et al., 2014), a probe created by modifying 6E10 antibody with PEG and ^64^Cu that distinguished Tg from control mice (McLean et al., 2012), and a probe developed from an ^124^I-labeled mAb158 against Aβ protofibrils (Magnusson et al., 2013). Still, none of these probes specifically target AβOs.

Previously, we described a molecular MRI probe that is targeted against AβOs (Viola et al., 2015). Based on the success of our initial MRI probe and the antibody-based probes being explored by others, it follows that AβO-specific antibodies can be used to target probes and provide better signal-to-noise ratios. Here we showed that anti-AβO antibodies can be used to develop molecular MRI ad PET probes that distinguish WT mice from their 5xFAD littermates at ages as early as 4 months old. These probes have proven to be non-toxic over the periods examined and, in fact, showed *in vivo* efficacy. These studies, however, are limited to the 5xFAD mouse model for AD and have not yet been tested in other animal models or in human subjects. Our paper in essence establishes proof of concept that oligomers can be detected by antibody-based probes for PET and MRI. This is a first step, and a great deal of work remains. A case in point, while *ex vivo* PET imaging is robust in its ability to distinguish AD from control brains, the conditions for *in vivo* imaging require significant optimization.

Early diagnostics are critical to combating this devastating disease, but without effective therapeutics, they have limited value. The first FDA-approved drug to treat Alzheimer’s disease (AD) in nearly two decades, Aduhelm^®^, shows a preferential affinity for all aggregated forms of amyloid beta (Aβ), rather than targeting only the toxic AβOs. Currently, there are more than 126 agents in clinical trials, with most aimed at disease modification (Cummings, 2021; Cummings et al., 2021). While less than 10% of these target Aβ, there remains evidence that Aβ is a significant target for therapeutic development. Lowering AβO levels by enhancing fibril formation has been shown to be protective (Mucke et al., 2000). This is supported by previous antibody-based studies (Lambert et al., 2007; Xiao et al., 2013). The data presented here importantly show that AβO-selective antibodies rescue memory performance in a widely used AD model. These antibodies, which have been modified for use in brain imaging of AβO, show great promise as potential agents for AD therapeutics and diagnostics; the potential of one AβO-selective antibody is now being assessed in a recently begun clinical trial.

## Abbreviations

AD: Alzheimer’s disease
AβO: Amyloid β oligomer
CSF: cerebrospinal fluid
GFAP: glial fibrillar acidic protein
ICV: intracerebroventricular
MNS: magnetic nanostructures
MRI: magnetic resonance imaging
PET: positron emission tomography
NOR: novel object recognition
NLR: novel location recognition
PiB: Pittsburgh Compound B
pTau: phosphorylated tau
ThioS: thioflavin S
Tg: transgenic
WT: wild-type

## Acknowledgements

We would like to thank Samuel C. Bartley, Elizabeth A. Johnson, Matthew Perkins, Jake Vitrofsky, Alex L. Qin, Henry Weiss, Rohan Chalasani, and Erika N. Cline for their assistance that helped make this study possible. We would like to thank the Northwestern University Research Experiences for Undergraduates program for their support.

This work was supported in part by NIH grants (R41AG054337, R21AG045637 to WLK and RF1AG063903 to Kelleher, Patrie, and WLK). E.A.W. is supported by grant number 2020-225578 from the Chan Zuckerberg Initiative DAF, an advised fund of Silicon Valley Community Foundation.

Microscopy was performed at the Biological Imaging Facility at Northwestern University (RRID:SCR_017767), graciously supported by the Chemistry for Life Processes Institute, the NU Office for Research, the Department of Molecular Biosciences and the Rice Foundation**.

MRI and PET/CT imaging work was performed at the Northwestern University Center for Advanced Molecular Imaging (RRID:SCR_021192), graciously supported by the Chemistry for Life Processes Institute, the NU Office for Research. Imaging generously supported by NCI CCSG P30 CA060553 awarded to the Robert H Lurie Comprehensive Cancer Center.

Imaging work was performed at the Northwestern University High-Throughput Analysis Lab, graciously supported by the Chemistry for Life Processes Institute, the NU Office for Research.

## Contribution to the field statement

Alzheimer’s disease is costly and marked by pathological damage and progressive memory loss. While there has been progress made towards developing better therapeutics and diagnostics, it has been limited. Diagnostic improvements have primarily been in the development of better imaging methods, mostly using agents that probe amyloid fibrils and plaques-species that do not correlate well with disease progression and are not present at the earliest stages of the disease. Amyloid β oligomers (AβOs) are now widely accepted as the Aβ species most germane to AD onset and progression. Here we report evidence further supporting the role of AβOs in Alzheimer’s disease and introduce a promising anti-AβO diagnostic probe capable of distinguishing the 5xFAD mouse model from wild type mice by PET and MRI. Our studies also showed a concomitant development of memory impairment with the accumulation of AβOs and reactive astrocytes. Compelling support for the conclusion that AβOs cause memory loss was found in experiments showing that AβO-selective antibodies into 5xFAD mice completely restored memory function. These antibodies, modified to give imaging probes, were able to distinguish 5xFAD mice from wild type littermates. These results demonstrate that AβO selective antibodies have potential both for therapeutics and for diagnostics.

## References

(2021). 2021 Alzheimer’s disease facts and figures. Alzheimers Dement 17(3), 327–406. doi: 10.1002/alz.12328.

Acton, P.Q., PA, US), An, Z.A., PA, US), Bett, A.J.L., PA, US), Breese, R.Q., PA, US), Chang, L.W., IL, US), Dodson, E.C.S., PA, US), et al. (2010). Anti-ADDL antibodies and uses thereof. United States patent application 11/256332. 08/24/2010.

Albert, M.S., DeKosky, S.T., Dickson, D., Dubois, B., Feldman, H.H., Fox, N.C., et al. (2011). The diagnosis of mild cognitive impairment due to Alzheimer’s disease: recommendations from the National Institute on Aging-Alzheimer’s Association workgroups on diagnostic guidelines for Alzheimer’s disease. Alzheimers Dement 7(3), 270–279. doi: 10.1016/j.jalz.2011.03.008.

Antunes, M., and Biala, G. (2012). The novel object recognition memory: neurobiology, test procedure, and its modifications. Cogn Process 13(2), 93–110. doi: 10.1007/s10339-011-0430-z.

Ashe, K.H. (2020). The biogenesis and biology of amyloid beta oligomers in the brain. Alzheimers Dement 16(11), 1561–1567. doi: 10.1002/alz.12084.

Ashok, A., Singh, N., Chaudhary, S., Bellamkonda, V., Kritikos, A.E., Wise, A.S., et al. (2020). Retinal Degeneration and Alzheimer’s Disease: An Evolving Link. Int J Mol Sci 21(19). doi: 10.3390/ijms21197290.

Bastin, C., Bahri, M.A., Meyer, F., Manard, M., Delhaye, E., Plenevaux, A., et al. (2020). In vivo imaging of synaptic loss in Alzheimer’s disease with [18F]UCB-H positron emission tomography. Eur J Nucl Med Mol Imaging 47(2), 390–402. doi: 10.1007/s00259-019-04461-x.

Bengoetxea, X., Rodriguez-Perdigon, M., and Ramirez, M.J. (2015). Object recognition test for studying cognitive impairments in animal models of Alzheimer’s disease. Front Biosci (Schol Ed) 7, 10–29.

Benitez, B.A., Jin, S.C., Guerreiro, R., Graham, R., Lord, J., Harold, D., et al. (2014). Missense variant in TREML2 protects against Alzheimer’s disease. Neurobiol Aging 35(6), 1510.e1519–1526. doi: 10.1016/j.neurobiolaging.2013.12.010.

Bicca, M.A., Costa, R., Loch-Neckel, G., Figueiredo, C.P., Medeiros, R., and Calixto, J.B. (2015). B(2) receptor blockage prevents Abeta-induced cognitive impairment by neuroinflammation inhibition. Behav Brain Res 278, 482–491. doi: 10.1016/j.bbr.2014.10.040.

Braak, H., and Del Tredici, K. (2011). Alzheimer’s pathogenesis: is there neuron-to-neuron propagation? Acta Neuropathol 121(5), 589–595. doi: 10.1007/s00401-011-0825-z.

Buchanan, H., Mackay, M., Palmer, K., Tothová, K., Katsur, M., Platt, B., et al. (2020). Synaptic Loss, ER Stress and Neuro-Inflammation Emerge Late in the Lateral Temporal Cortex and Associate with Progressive Tau Pathology in Alzheimer’s Disease. Mol Neurobiol 57(8), 3258–3272. doi: 10.1007/s12035-020-01950-1.

Camporesi, E., Nilsson, J., Brinkmalm, A., Becker, B., Ashton, N.J., Blennow, K., et al. (2020). Fluid Biomarkers for Synaptic Dysfunction and Loss. Biomark Insights 15, 1177271920950319. doi: 10.1177/1177271920950319.

Chang, L., Bakhos, L., Wang, Z., Venton, D.L., and Klein, W.L. (2003). Femtomole immunodetection of synthetic and endogenous amyloid-beta oligomers and its application to Alzheimer’s disease drug candidate screening. J Mol Neurosci 20(3), 305–313. doi: 10.1385/JMN:20:3:305.

Cheng, Y.J., Lin, C.H., and Lane, H.Y. (2021). From Menopause to Neurodegeneration-Molecular Basis and Potential Therapy. Int J Mol Sci 22(16). doi: 10.3390/ijms22168654.

Choi, M., Kim, H., Yang, E.J., and Kim, H.S. (2020). Inhibition of STAT3 phosphorylation attenuates impairments in learning and memory in 5XFAD mice, an animal model of Alzheimer’s disease. J Pharmacol Sci 143(4), 290–299. doi: 10.1016/j.jphs.2020.05.009.

Cline, E., Viola, K., Klein, W., Wang, X., Bacskai, B., Rammes, G., et al. (2019a). “Synaptic intervention in Alzheimer’s disease: soluble Aβ oligomer directed ACU193 monoclonal antibody therapeutic for treatment of early Alzheimer’s disease”, in: Clinical Trials on Alzheimer’s disease. (San Diego, CA, USA).

Cline, E.N., Bicca, M.A., Viola, K.L., and Klein, W.L. (2018). The Amyloid-beta Oligomer Hypothesis: Beginning of the Third Decade. J Alzheimers Dis 64(s1), S567–S610. doi: 10.3233/JAD-179941.

Cline, E.N., Das, A., Bicca, M.A., Mohammad, S.N., Schachner, L.F., Kamel, J.M., et al. (2019b). A novel crosslinking protocol stabilizes amyloid beta oligomers capable of inducing Alzheimer’s-associated pathologies. J Neurochem 148(6), 822–836. doi: 10.1111/jnc.14647.

Cohen, S.J., and Stackman, R.W., Jr. (2015). Assessing rodent hippocampal involvement in the novel object recognition task. A review. Behav Brain Res 285, 105–117. doi: 10.1016/j.bbr.2014.08.002.

Colom-Cadena, M., Spires-Jones, T., Zetterberg, H., Blennow, K., Caggiano, A., DeKosky, S.T., et al. (2020). The clinical promise of biomarkers of synapse damage or loss in Alzheimer’s disease. Alzheimers Res Ther 12(1), 21. doi: 10.1186/s13195-020-00588-4.

Cummings, J. (2021). Drug Development for Psychotropic, Cognitive-Enhancing, and Disease-Modifying Treatments for Alzheimer’s Disease. J Neuropsychiatry Clin Neurosci 33(1), 3–13. doi: 10.1176/appi.neuropsych.20060152.

Cummings, J., Lee, G., Zhong, K., Fonseca, J., and Taghva, K. (2021). Alzheimer’s disease drug development pipeline: 2021. Alzheimers Dement (N Y) 7(1), e12179. doi: 10.1002/trc2.12179.

De Felice, F.G., Vieira, M.N., Bomfim, T.R., Decker, H., Velasco, P.T., Lambert, M.P., et al. (2009). Protection of synapses against Alzheimer’s-linked toxins: insulin signaling prevents the pathogenic binding of Abeta oligomers. Proc Natl Acad Sci U S A 106(6), 1971–1976. doi: 10.1073/pnas.0809158106.

Denninger, J.K., Smith, B.M., and Kirby, E.D. (2018). Novel Object Recognition and Object Location Behavioral Testing in Mice on a Budget. J Vis Exp (141). doi: 10.3791/58593.

Devi, L., Alldred, M.J., Ginsberg, S.D., and Ohno, M. (2010). Sex- and brain region-specific acceleration of beta-amyloidogenesis following behavioral stress in a mouse model of Alzheimer’s disease. Mol Brain 3, 34. doi: 10.1186/1756-6606-3-34.

Esparza, T.J., Zhao, H., Cirrito, J.R., Cairns, N.J., Bateman, R.J., Holtzman, D.M., et al. (2013). Amyloid-beta oligomerization in Alzheimer dementia versus high-pathology controls. Ann Neurol 73(1), 104–119. doi: 10.1002/ana.23748.

Ferreira, S.T., and Klein, W.L. (2011). The Abeta oligomer hypothesis for synapse failure and memory loss in Alzheimer’s disease. Neurobiol Learn Mem 96(4), 529–543. doi: 10.1016/j.nlm.2011.08.003.

Gandy, S., Simon, A.J., Steele, J.W., Lublin, A.L., Lah, J.J., Walker, L.C., et al. (2010). Days to criterion as an indicator of toxicity associated with human Alzheimer amyloid-beta oligomers. Ann Neurol 68(2), 220–230. doi: 10.1002/ana.22052.

Georganopoulou, D.G., Chang, L., Nam, J.M., Thaxton, C.S., Mufson, E.J., Klein, W.L., et al. (2005). Nanoparticle-based detection in cerebral spinal fluid of a soluble pathogenic biomarker for Alzheimer’s disease. Proc Natl Acad Sci U S A 102(7), 2273–2276. doi: 10.1073/pnas.0409336102.

Girard, S.D., Baranger, K., Gauthier, C., Jacquet, M., Bernard, A., Escoffier, G., et al. (2013). Evidence for early cognitive impairment related to frontal cortex in the 5XFAD mouse model of Alzheimer’s disease. J Alzheimers Dis 33(3), 781–796. doi: 10.3233/jad-2012-120982.

Girard, S.D., Jacquet, M., Baranger, K., Migliorati, M., Escoffier, G., Bernard, A., et al. (2014). Onset of hippocampus-dependent memory impairments in 5XFAD transgenic mouse model of Alzheimer’s disease. Hippocampus 24(7), 762–772. doi: 10.1002/hipo.22267.

Gong, Y., Chang, L., Viola, K.L., Lacor, P.N., Lambert, M.P., Finch, C.E., et al. (2003). Alzheimer’s disease-affected brain: presence of oligomeric A beta ligands (ADDLs) suggests a molecular basis for reversible memory loss. Proc Natl Acad Sci U S A 100(18), 10417–10422. doi: 10.1073/pnas.1834302100.

Goure, W.F., Krafft, G.A., Jerecic, J., and Hefti, F. (2014). Targeting the proper amyloid-beta neuronal toxins: a path forward for Alzheimer’s disease immunotherapeutics. Alzheimers Res Ther 6(4), 42. doi: 10.1186/alzrt272.

Grayson, B., Leger, M., Piercy, C., Adamson, L., Harte, M., and Neill, J.C. (2015). Assessment of disease-related cognitive impairments using the novel object recognition (NOR) task in rodents. Behav Brain Res 285, 176–193. doi: 10.1016/j.bbr.2014.10.025.

Guntern, R., Bouras, C., Hof, P.R., and Vallet, P.G. (1992). An improved thioflavine S method for staining neurofibrillary tangles and senile plaques in Alzheimer’s disease. Experientia 48(1), 8–10. doi: 10.1007/BF01923594.

Guven, G., Bilgic, B., Samanci, B., Gurvit, H., Hanagasi, H., Donmez, C., et al. (2020). Peripheral TREM2 mRNA levels in early and late-onset Alzheimer disease’s patients. Mol Biol Rep 47(8), 5903–5909. doi: 10.1007/s11033-020-05661-7.

Guzman-Martinez, L., Maccioni, R.B., Farías, G.A., Fuentes, P., and Navarrete, L.P. (2019). Biomarkers for Alzheimer’s Disease. Curr Alzheimer Res 16(6), 518–528. doi: 10.2174/1567205016666190517121140.

Hampel, H., Hardy, J., Blennow, K., Chen, C., Perry, G., Kim, S.H., et al. (2021). The Amyloid-beta Pathway in Alzheimer’s Disease. Mol Psychiatry. doi: 10.1038/s41380-021-01249-0.

Hefti, F., Goure, W.F., Jerecic, J., Iverson, K.S., Walicke, P.A., and Krafft, G.A. (2013). The case for soluble Abeta oligomers as a drug target in Alzheimer’s disease. Trends Pharmacol Sci 34(5), 261–266. doi: 10.1016/j.tips.2013.03.002.

Hsia, A.Y., Masliah, E., McConlogue, L., Yu, G.Q., Tatsuno, G., Hu, K., et al. (1999). Plaque-independent disruption of neural circuits in Alzheimer’s disease mouse models. Proc Natl Acad Sci U S A 96(6), 3228–3233.

Huang, S., Tong, H., Lei, M., Zhou, M., Guo, W., Li, G., et al. (2017). Astrocytic glutamatergic transporters are involved in Abeta-induced synaptic dysfunction. Brain Res. doi: 10.1016/j.brainres.2017.10.011.

Hulbert, S., and Adeli, H. (2013). EEG/MEG- and imaging-based diagnosis of Alzheimer’s disease. Rev Neurosci 24(6), 563–576. doi: 10.1515/revneuro-2013-0042.

Investor Relations, B. (2021). “FDA grants accelerated approval for ADUHELM™ as the first and only Alzheimer’s disease treatment to address a defining pathology of the disease”. (www.biogen.com:Biogen).

Jack, C.R., Jr., Albert, M.S., Knopman, D.S., McKhann, G.M., Sperling, R.A., Carrillo, M.C., et al. (2011). Introduction to the recommendations from the National Institute on Aging-Alzheimer’s Association workgroups on diagnostic guidelines for Alzheimer’s disease. Alzheimers Dement 7(3), 257–262. doi: 10.1016/j.jalz.2011.03.004.

Jawhar, S., Trawicka, A., Jenneckens, C., Bayer, T.A., and Wirths, O. (2012). Motor deficits, neuron loss, and reduced anxiety coinciding with axonal degeneration and intraneuronal Abeta aggregation in the 5XFAD mouse model of Alzheimer’s disease. Neurobiol Aging 33(1), 196 e129–140. doi: 10.1016/j.neurobiolaging.2010.05.027.

Jiang, T., Yu, J.T., Zhu, X.C., and Tan, L. (2013). TREM2 in Alzheimer’s disease. Mol Neurobiol 48(1), 180–185. doi: 10.1007/s12035-013-8424-8.

Johnson, K.A., Minoshima, S., Bohnen, N.I., Donohoe, K.J., Foster, N.L., Herscovitch, P., et al. (2013). Appropriate use criteria for amyloid PET: a report of the Amyloid Imaging Task Force, the Society of Nuclear Medicine and Molecular Imaging, and the Alzheimer’s Association. Alzheimers Dement 9(1), e-1–16. doi: 10.1016/j.jalz.2013.01.002.

Kanno, T., Tsuchiya, A., and Nishizaki, T. (2014). Hyperphosphorylation of Tau at Ser396 occurs in the much earlier stage than appearance of learning and memory disorders in 5XFAD mice. Behav Brain Res 274, 302–306. doi: 10.1016/j.bbr.2014.08.034.

Kayed, R., Head, E., Thompson, J.L., McIntire, T.M., Milton, S.C., Cotman, C.W., et al. (2003). Common structure of soluble amyloid oligomers implies common mechanism of pathogenesis. Science 300(5618), 486–489. doi: 10.1126/science.1079469.

Kimura, R., and Ohno, M. (2009). Impairments in remote memory stabilization precede hippocampal synaptic and cognitive failures in 5XFAD Alzheimer mouse model. Neurobiol Dis 33(2), 229–235. doi: 10.1016/j.nbd.2008.10.006.

Klunk, W.E., Engler, H., Nordberg, A., Wang, Y., Blomqvist, G., Holt, D.P., et al. (2004). Imaging brain amyloid in Alzheimer’s disease with Pittsburgh Compound-B. Ann Neurol 55(3), 306–319. doi: 10.1002/ana.20009.

Krafft, G., Hefti, F., Goure, W., Jerecic, J., Iverson, K., and Walicke, P. (2013). ACU-193: A candidate therapeutic antibody that selectively targets soluble beta-amyloid oligomers. Alzheimer’s & Dementia 9(4, Supplement), P326. doi: http://dx.doi.orq/10.1016/j.jalz.2013.04.166.

Lacor, P.N., Buniel, M.C., Chang, L., Fernandez, S.J., Gong, Y., Viola, K.L., et al. (2004). Synaptic targeting by Alzheimer’s-related amyloid beta oligomers. J Neurosci 24(45), 10191–10200. doi: 10.1523/JNEUROSCI.3432-04.2004.

Lacor, P.N., Buniel, M.C., Furlow, P.W., Clemente, A.S., Velasco, P.T., Wood, M., et al. (2007). Abeta oligomer-induced aberrations in synapse composition, shape, and density provide a molecular basis for loss of connectivity in Alzheimer’s disease. J Neurosci 27(4), 796–807. doi: 10.1523/JNEUROSCI.3501-06.2007.

Lambert, M.P., Barlow, A.K., Chromy, B.A., Edwards, C., Freed, R., Liosatos, M., et al. (1998). Diffusible, nonfibrillar ligands derived from Abeta1-42 are potent central nervous system neurotoxins. Proc Natl Acad Sci U S A 95(11), 6448–6453.

Lambert, M.P., Velasco, P.T., Chang, L., Viola, K.L., Fernandez, S., Lacor, P.N., et al. (2007). Monoclonal antibodies that target pathological assemblies of Abeta. J Neurochem 100(1), 23–35. doi: 10.1111/j.1471-4159.2006.04157.x.

Lasagna-Reeves, C.A., Castillo-Carranza, D.L., Sengupta, U., Guerrero-Munoz, M.J., Kiritoshi, T., Neugebauer, V., et al. (2012). Alzheimer brain-derived tau oligomers propagate pathology from endogenous tau. Sci Rep 2, 700. doi: 10.1038/srep00700.

Lee, S.P., Falangola, M.F., Nixon, R.A., Duff, K., and Helpern, J.A. (2004). Visualization of beta-amyloid plaques in a transgenic mouse model of Alzheimer’s disease using MR microscopy without contrast reagents. Magn Reson Med 52(3), 538–544. doi: 10.1002/mrm.20196.

Lesne, S., Koh, M.T., Kotilinek, L., Kayed, R., Glabe, C.G., Yang, A., et al. (2006). A specific amyloid-beta protein assembly in the brain impairs memory. Nature 440(7082), 352–357. doi: 10.1038/nature04533.

Li, S., and Selkoe, D.J. (2020). A mechanistic hypothesis for the impairment of synaptic plasticity by soluble Abeta oligomers from Alzheimer’s brain. J Neurochem 154(6), 583–597. doi: 10.1111/jnc.15007.

Lin, N., Gao, J., Mao, C., Sun, H., Lu, Q., and Cui, L. (2021). Differences in Multimodal Electroencephalogram and Clinical Correlations Between Early-Onset Alzheimer’s Disease and Frontotemporal Dementia. Front Neurosci 15, 687053. doi: 10.3389/fnins.2021.687053.

Magnusson, K., Sehlin, D., Syvanen, S., Svedberg, M.M., Philipson, O., Soderberg, L., et al. (2013). Specific uptake of an amyloid-beta protofibril-binding antibody-tracer in AbetaPP transgenic mouse brain. J Alzheimers Dis 37(1), 29–40. doi: 10.3233/jad-130029.

Masters, C.L., Simms, G., Weinman, N.A., Multhaup, G., McDonald, B.L., and Beyreuther, K. (1985). Amyloid plaque core protein in Alzheimer disease and Down syndrome. Proc Natl Acad Sci U S A 82(12), 4245–4249.

Mathis, C.A., Wang, Y., Holt, D.P., Huang, G.F., Debnath, M.L., and Klunk, W.E. (2003). Synthesis and evaluation of 11C-labeled 6-substituted 2-arylbenzothiazoles as amyloid imaging agents. J Med Chem 46(13), 2740–2754. doi: 10.1021/jm030026b.

McKhann, G.M., Knopman, D.S., Chertkow, H., Hyman, B.T., Jack, C.R., Jr., Kawas, C.H., et al. (2011). The diagnosis of dementia due to Alzheimer’s disease: recommendations from the National Institute on Aging-Alzheimer’s Association workgroups on diagnostic guidelines for Alzheimer’s disease. Alzheimers Dement 7(3), 263–269. doi: 10.1016/j.jalz.2011.03.005.

McLean, D., Cooke, M.J., Wang, Y., Green, D., Fraser, P.E., George-Hyslop, P.S., et al. (2012). Anti-amyloid-beta-mediated positron emission tomography imaging in Alzheimer’s disease mouse brains. PLoS One 7(12), e51958. doi: 10.1371/journal.pone.0051958.

Mecca, A.P., Chen, M.K., O’Dell, R.S., Naganawa, M., Toyonaga, T., Godek, T.A., et al. (2020). In vivo measurement of widespread synaptic loss in Alzheimer’s disease with SV2A PET. Alzheimers Dement 16(7), 974–982. doi: 10.1002/alz.12097.

Meng, X., Li, T., Wang, X., Lv, X., Sun, Z., Zhang, J., et al. (2019). Association between increased levels of amyloid-β oligomers in plasma and episodic memory loss in Alzheimer’s disease. Alzheimer’s Research & Therapy 11(1), 89. doi: 10.1186/s13195-019-0535-7.

Mirzaei, N., Shi, H., Oviatt, M., Doustar, J., Rentsendorj, A., Fuchs, D.T., et al. (2020). Alzheimer’s Retinopathy: Seeing Disease in the Eyes. Front Neurosci 14, 921. doi: 10.3389/fnins.2020.00921.

Monterey, M.D., Wei, H., Wu, X., and Wu, J.Q. (2021). The Many Faces of Astrocytes in Alzheimer’s Disease. Front Neurol 12, 619626. doi: 10.3389/fneur.2021.619626.

Montoliu-Gaya, L., Strydom, A., Blennow, K., Zetterberg, H., and Ashton, N.J. (2021). Blood Biomarkers for Alzheimer’s Disease in Down Syndrome. J Clin Med 10(16). doi: 10.3390/jcm10163639.

Mucke, L., Masliah, E., Yu, G.Q., Mallory, M., Rockenstein, E.M., Tatsuno, G., et al. (2000). High-level neuronal expression of abeta 1-42 in wild-type human amyloid protein precursor transgenic mice: synaptotoxicity without plaque formation. Journal of Neuroscience 20(11), 4050–4058.

Mucke, L., and Selkoe, D.J. (2012). Neurotoxicity of amyloid beta-protein: synaptic and network dysfunction. Cold Spring Harb Perspect Med 2(7), a006338. doi: 10.1101/cshperspect.a006338.

Mufson, E.J., Ward, S., and Binder, L. (2014). Prefibrillar tau oligomers in mild cognitive impairment and Alzheimer’s disease. Neurodegener Dis 13(2-3), 151–153. doi: 10.1159/000353687.

Mundt, A.P., Winter, C., Mueller, S., Wuerfel, J., Tysiak, E., Schnorr, J., et al. (2009). Targeting activated microglia in Alzheimer’s pathology by intraventricular delivery of a phagocytosable MRI contrast agent in APP23 transgenic mice. Neuroimage 46(2), 367–372. doi: 10.1016/j.neuroimage.2009.01.067.

Nandwana, V., Ryoo, S.-R., Kanthala, S., De, M., Chou, S.S., Prasad, P.V., et al. (2016). Engineered Theranostic Magnetic Nanostructures: Role of Composition and Surface Coating on Magnetic Resonance Imaging Contrast and Thermal Activation. ACS Applied Materials & Interfaces 8(11), 6953–6961. doi: 10.1021/acsami.6b01377.

Nisa, F.Y., Rahman, M.A., Hossen, M.A., Khan, M.F., Khan, M.A.N., Majid, M., et al. (2021). Role of neurotoxicants in the pathogenesis of Alzheimer’s disease: a mechanistic insight. Ann Med 53(1), 1476–1501. doi: 10.1080/07853890.2021.1966088.

Nyborg, A.C., Moll, J.R., Wegrzyn, R.D., Havas, D., Hutter-Paier, B., Feuerstein, G.G., et al. (2013). In vivo and ex vivo imaging of amyloid-beta cascade aggregates with a Pronucleon peptide. J Alzheimers Dis 34(4), 957–967. doi: 10.3233/jad-122107.

Oakley, H., Cole, S.L., Logan, S., Maus, E., Shao, P., Craft, J., et al. (2006). Intraneuronal beta-amyloid aggregates, neurodegeneration, and neuron loss in transgenic mice with five familial Alzheimer’s disease mutations: potential factors in amyloid plaque formation. J Neurosci 26(40), 10129–10140. doi: 10.1523/JNEUROSCI.1202-06.2006.

Oblak, A.L., Lin, P.B., Kotredes, K.P., Pandey, R.S., Garceau, D., Williams, H.M., et al. (2021). Comprehensive Evaluation of the 5XFAD Mouse Model for Preclinical Testing Applications: A MODEL-AD Study. Front Aging Neurosci 13, 713726. doi: 10.3389/fnagi.2021.713726.

Ohno, M. (2009). Failures to reconsolidate memory in a mouse model of Alzheimer’s disease. Neurobiol Learn Mem 92(3), 455–459. doi: 10.1016/j.nlm.2009.05.001.

Ohno, M., Chang, L., Tseng, W., Oakley, H., Citron, M., Klein, W.L., et al. (2006). Temporal memory deficits in Alzheimer’s mouse models: rescue by genetic deletion of BACE1. Eur J Neurosci 23(1), 251–260. doi: 10.1111/j.1460-9568.2005.04551.x.

Ou-Yang, M.H., and Van Nostrand, W.E. (2013). The absence of myelin basic protein promotes neuroinflammation and reduces amyloid beta-protein accumulation in Tg-5xFAD mice. J Neuroinflammation 10, 134. doi: 10.1186/1742-2094-10-134.

Overk, C.R., and Masliah, E. (2014). Toward a unified therapeutics approach targeting putative amyloid-beta oligomer receptors. Proc Natl Acad Sci U S A 111(38), 13680–13681. doi: 10.1073/pnas.1414554111.

Park, S.Y., Avraham, H.K., and Avraham, S. (2004). RAFTK/Pyk2 activation is mediated by trans-acting autophosphorylation in a Src-independent manner. J Biol Chem 279(32), 33315–33322. doi: 10.1074/jbc.M313527200.

Pereira, J.B., Janelidze, S., Ossenkoppele, R., Kvartsberg, H., Brinkmalm, A., Mattsson-Carlgren, N., et al. (2021). Untangling the association of amyloid-β and tau with synaptic and axonal loss in Alzheimer’s disease. Brain 144(1), 310–324. doi: 10.1093/brain/awaa395.

Pitt, J., Wilcox, K.C., Tortelli, V., Diniz, L.P., Oliveira, M.S., Dobbins, C., et al. (2017). Neuroprotective astrocyte-derived insulin/insulin-like growth factor 1 stimulates endocytic processing and extracellular release of neuron-bound Abeta oligomers. Mol Biol Cell 28(20), 2623–2636. doi: 10.1091/mbc.E17-06-0416.

Preeti, K., Sood, A., and Fernandes, V. (2021). Metabolic Regulation of Glia and Their Neuroinflammatory Role in Alzheimer’s Disease. Cell Mol Neurobiol. doi: 10.1007/s10571-021-01147-7.

Price, B.R., Sudduth, T.L., Weekman, E.M., Johnson, S., Hawthorne, D., Woolums, A., et al. (2020). Therapeutic Trem2 activation ameliorates amyloid-beta deposition and improves cognition in the 5XFAD model of amyloid deposition. J Neuroinflammation 17(1), 238. doi: 10.1186/s12974-020-01915-0.

Robakis, N.K. (2011). Mechanisms of AD neurodegeneration may be independent of Abeta and its derivatives. Neurobiol Aging 32(3), 372–379. doi: 10.1016/j.neurobiolaging.2010.05.022.

Rodgers, A.B. (2005). “Progress report on Alzheimer’s disease 2004-2005”. U.S.Department of Health and Human Services; National Institutes on Aging; National Institutes of Health).

Rokka, J., Snellman, A., Zona, C., La Ferla, B., Nicotra, F., Salmona, M., et al. (2014). Synthesis and evaluation of a (18)F-curcumin derivate for beta-amyloid plaque imaging. Bioorg Med Chem 22(9), 2753–2762. doi: 10.1016/j.bmc.2014.03.010.

Savage, M.J., Kalinina, J., Wolfe, A., Tugusheva, K., Korn, R., Cash-Mason, T., et al. (2014). A sensitive abeta oligomer assay discriminates Alzheimer’s and aged control cerebrospinal fluid. J Neurosci 34(8), 2884–2897. doi: 10.1523/jneurosci.1675-13.2014.

Schnabel, J. (2011). Amyloid: little proteins, big clues. Nature 475(7355), S12–14. doi: 10.1038/475S12a.

Selkoe, D.J., and Hardy, J. (2016). The amyloid hypothesis of Alzheimer’s disease at 25 years. EMBO Mol Med 8(6), 595–608. doi: 10.15252/emmm.201606210.

Shao, C.Y., Mirra, S.S., Sait, H.B., Sacktor, T.C., and Sigurdsson, E.M. (2011). Postsynaptic degeneration as revealed by PSD-95 reduction occurs after advanced Abeta and tau pathology in transgenic mouse models of Alzheimer’s disease. Acta Neuropathol 122(3), 285–292. doi: 10.1007/s00401-011-0843-x.

Siwek, M.E., Muller, R., Henseler, C., Trog, A., Lundt, A., Wormuth, C., et al. (2015). Altered Theta Oscillations and Aberrant Cortical Excitatory Activity in the 5XFAD Model of Alzheimer’s Disease. Neural Plast 2015, 781731. doi: 10.1155/2015/781731.

Slemmon, J.R., Meredith, J., Guss, V., Andreasson, U., Andreasen, N., Zetterberg, H., et al. (2012). Measurement of Abeta1-42 in cerebrospinal fluid is influenced by matrix effects. J Neurochem 120(2), 325–333. doi: 10.1111/j.1471-4159.2011.07553.x.

Sperling, R.A., Aisen, P.S., Beckett, L.A., Bennett, D.A., Craft, S., Fagan, A.M., et al. (2011). Toward defining the preclinical stages of Alzheimer’s disease: recommendations from the National Institute on Aging-Alzheimer’s Association workgroups on diagnostic guidelines for Alzheimer’s disease. Alzheimers Dement 7(3), 280–292. doi: 10.1016/j.jalz.2011.03.003.

Terry, R.D., Masliah, E., Salmon, D.P., Butters, N., DeTeresa, R., Hill, R., et al. (1991). Physical basis of cognitive alterations in Alzheimer’s disease: synapse loss is the major correlate of cognitive impairment. Ann Neurol 30(4), 572–580. doi: 10.1002/ana.410300410.

Toledo, J.B., Korff, A., Shaw, L.M., Trojanowski, J.Q., and Zhang, J. (2013a). CSF alpha-synuclein improves diagnostic and prognostic performance of CSF tau and Abeta in Alzheimer’s disease. Acta Neuropathol 126(5), 683–697. doi: 10.1007/s00401-013-1148-z.

Toledo, J.B., Xie, S.X., Trojanowski, J.Q., and Shaw, L.M. (2013b). Longitudinal change in CSF Tau and Abeta biomarkers for up to 48 months in ADNI. Acta Neuropathol 126(5), 659–670. doi: 10.1007/s00401-013-1151-4.

Townsend, M., Shankar, G.M., Mehta, T., Walsh, D.M., and Selkoe, D.J. (2006). Effects of secreted oligomers of amyloid beta-protein on hippocampal synaptic plasticity: a potent role for trimers. J Physiol 572(Pt 2), 477–492. doi: 10.1113/jphysiol.2005.103754.

Velasco, P.T., Heffern, M.C., Sebollela, A., Popova, I.A., Lacor, P.N., Lee, K.B., et al. (2012). Synapse-binding subpopulations of Abeta oligomers sensitive to peptide assembly blockers and scFv antibodies. ACS Chem Neurosci 3(11), 972–981. doi: 10.1021/cn300122k.

Viola, K.L., and Klein, W.L. (2015). Amyloid beta oligomers in Alzheimer’s disease pathogenesis, treatment, and diagnosis. Acta Neuropathol 129(2), 183–206. doi: 10.1007/s00401-015-1386-3.

Viola, K.L., Sbarboro, J., Sureka, R., De, M., Bicca, M.A., Wang, J., et al. (2015). Towards non-invasive diagnostic imaging of early-stage Alzheimer’s disease. Nat Nanotechnol 10(1), 91–98. doi: 10.1038/nnano.2014.254.

Wang, C., Xiong, M., Gratuze, M., Bao, X., Shi, Y., Andhey, P.S., et al. (2021). Selective removal of astrocytic APOE4 strongly protects against tau-mediated neurodegeneration and decreases synaptic phagocytosis by microglia. Neuron 109(10), 1657–1674 e1657. doi: 10.1016/j.neuron.2021.03.024.

Wang, H.W., Pasternak, J.F., Kuo, H., Ristic, H., Lambert, M.P., Chromy, B., et al. (2002). Soluble oligomers of beta amyloid (1-42) inhibit long-term potentiation but not long-term depression in rat dentate gyrus. Brain Res 924(2), 133–140.

Wipke, B.T., Wang, Z., Kim, J., McCarthy, T.J., and Allen, P.M. (2002). Dynamic visualization of a joint-specific autoimmune response through positron emission tomography. Nat Immunol 3(4), 366–372. doi: 10.1038/ni775.

Xiao, C., Davis, F.J., Chauhan, B.C., Viola, K.L., Lacor, P.N., Velasco, P.T., et al. (2013). Brain transit and ameliorative effects of intranasally delivered anti-amyloid-beta oligomer antibody in 5XFAD mice. J Alzheimers Dis 35(4), 777–788. doi: 10.3233/JAD-122419.

Yang, T., Dang, Y., Ostaszewski, B., Mengel, D., Steffen, V., Rabe, C., et al. (2019). Target engagement in an alzheimer trial: Crenezumab lowers amyloid beta oligomers in cerebrospinal fluid. Ann Neurol 86(2), 215–224. doi: 10.1002/ana.25513.

Yang, Y., Kim, J., Kim, H.Y., Ryoo, N., Lee, S., Kim, Y., et al. (2015). Amyloid-beta Oligomers May Impair SNARE-Mediated Exocytosis by Direct Binding to Syntaxin 1a. Cell Rep 12(8), 1244–1251. doi: 10.1016/j.celrep.2015.07.044.

Yuan, Q., Liu, X., Zhang, Y., Xian, Y.F., Zou, J., Zhang, X., et al. (2021). Established Beta Amyloid Pathology Is Unaffected by TREM2 Elevation in Reactive Microglia in an Alzheimer’s Disease Mouse Model. Molecules 26(9). doi: 10.3390/molecules26092685.

Zhang, M., Zhong, L., Han, X., Xiong, G., Xu, D., Zhang, S., et al. (2021a). Brain and Retinal Abnormalities in the 5xFAD Mouse Model of Alzheimer’s Disease at Early Stages. Front Neurosci 15, 681831. doi: 10.3389/fnins.2021.681831.

Zhang, T., Liao, Q., Zhang, D., Zhang, C., Yan, J., Ngetich, R., et al. (2021b). Predicting MCI to AD Conversation Using Integrated sMRI and rs-fMRI: Machine Learning and Graph Theory Approach. Front Aging Neurosci 13, 688926. doi: 10.3389/fnagi.2021.688926.

Zhao, Y., Wu, X., Li, X., Jiang, L.L., Gui, X., Liu, Y., et al. (2018). TREM2 Is a Receptor for β-Amyloid that Mediates Microglial Function. Neuron 97(5), 1023–1031.e1027. doi: 10.1016/j.neuron.2018.01.031.

Zhong, L., Xu, Y., Zhuo, R., Wang, T., Wang, K., Huang, R., et al. (2019). Soluble TREM2 ameliorates pathological phenotypes by modulating microglial functions in an Alzheimer’s disease model. Nat Commun 10(1), 1365. doi: 10.1038/s41467-019-09118-9.

Zhou, R., Ji, B., Kong, Y., Qin, L., Ren, W., Guan, Y., et al. (2021). PET Imaging of Neuroinflammation in Alzheimer’s Disease. Front Immunol 12, 739130. doi: 10.3389/fimmu.2021.739130.

