## Supplemental Figures for "The therapeutic and diagnostic potential of amyloid β oligomers (AβOs) selective antibodies to treat Alzheimer’s disease"

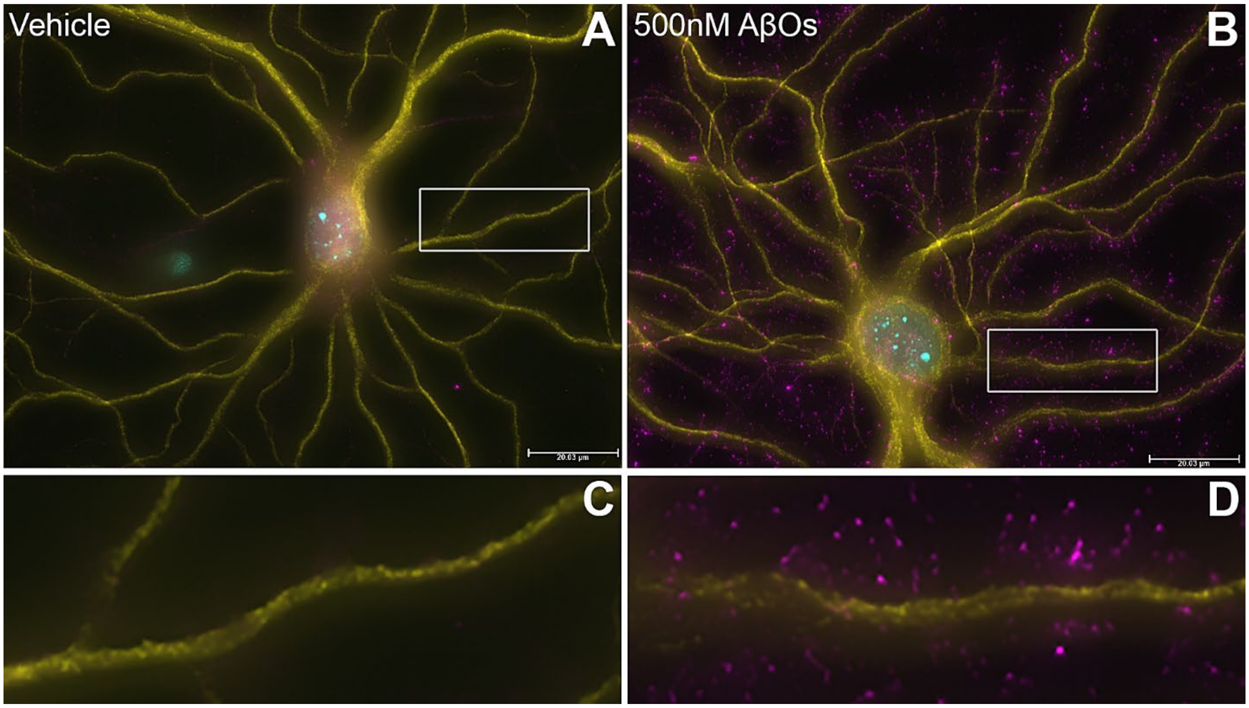


**Supplemental Figure 1**

**ACU193 detects AβOs bound to the spines of primary hippocampal neurons.** Primary hippocampal neurons treated with 500nM cross-linked AβOs (B, D) or volume equivalent of vehicle control (A, C) for 4 hours. Bound AβOs were detected using ACU193 antibody and imaged at 63x on the Leica DM6B fluorescent microscope. Images show robust, punctate labeling of AβOs (magenta) along the neuronal processes labeled with MAP2 (yellow). Magnified images (C, D) of processes show that the AβOs bind along spindly filaments projecting from the process (D), much like what has been shown by our lab previously (Lacor et al., 2004; Lacor et al., 2007; Pitt et al., 2017). Data show that, even with 10 times the amount of antibody present, almost no non-specific labeling was seen in the vehicle-treated cells.

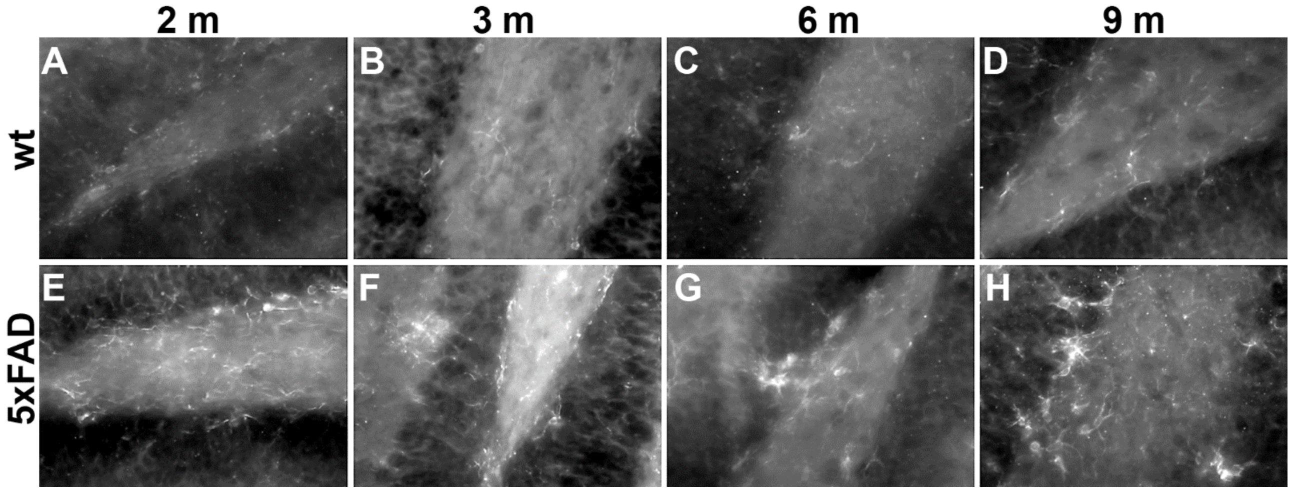


**Supplemental Figure 2**

**Microglia are activated by 2 months in 5xFAD mice.** (A-D) Wt mice show low levels of IbA1 immunoreactivity at all ages examined (2-9 months). In 5xFAD mice (E-H), IbA1 is elevated at all ages examined (2-9 months), with little change detected as AβO levels increase. Data would suggest that while GFAP increases concomitantly with AβO levels, IbA1 does not.

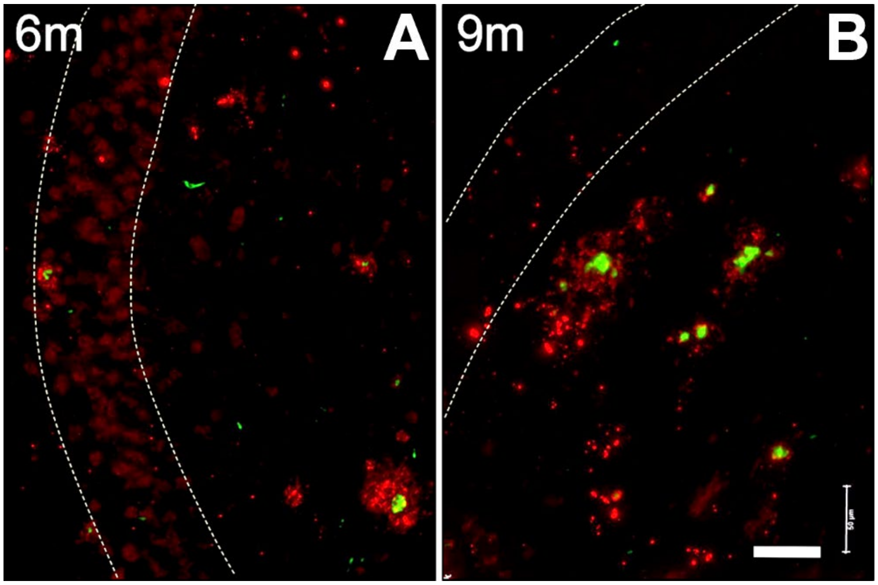


**Supplemental Figure 3**

**Immunohistochemistry of ACUDOTA distribution shows no overlap with ThioS stained plaques.** Brain sections from the hippocampus of 5xFAD mice (2-9 months) and WT littermates (9 month) were probed for AβO pathology using the humanized monoclonal ACU193 conjugated to DOTA (ACUDOTA), which is the non-radioactive form of ACUPET. Sections were counterstained with ThioS to detect dense core plaques. Dotted white lines indicate pyramidal cell layer. AβO pathology recognized by ACUDOTA (red) is detectable in proximity to ThioS-positive sites (green), but the labeling is distinct and does not co-localize. Immunohistochemistry shows that the PET probe based on the oligomer-selective ACU193 mAb does not bind plaques. Wildtype littermates showed neither plaques nor AβOs. Scale bar = 50μm. Data are representative of 6 samples per age and genotype.

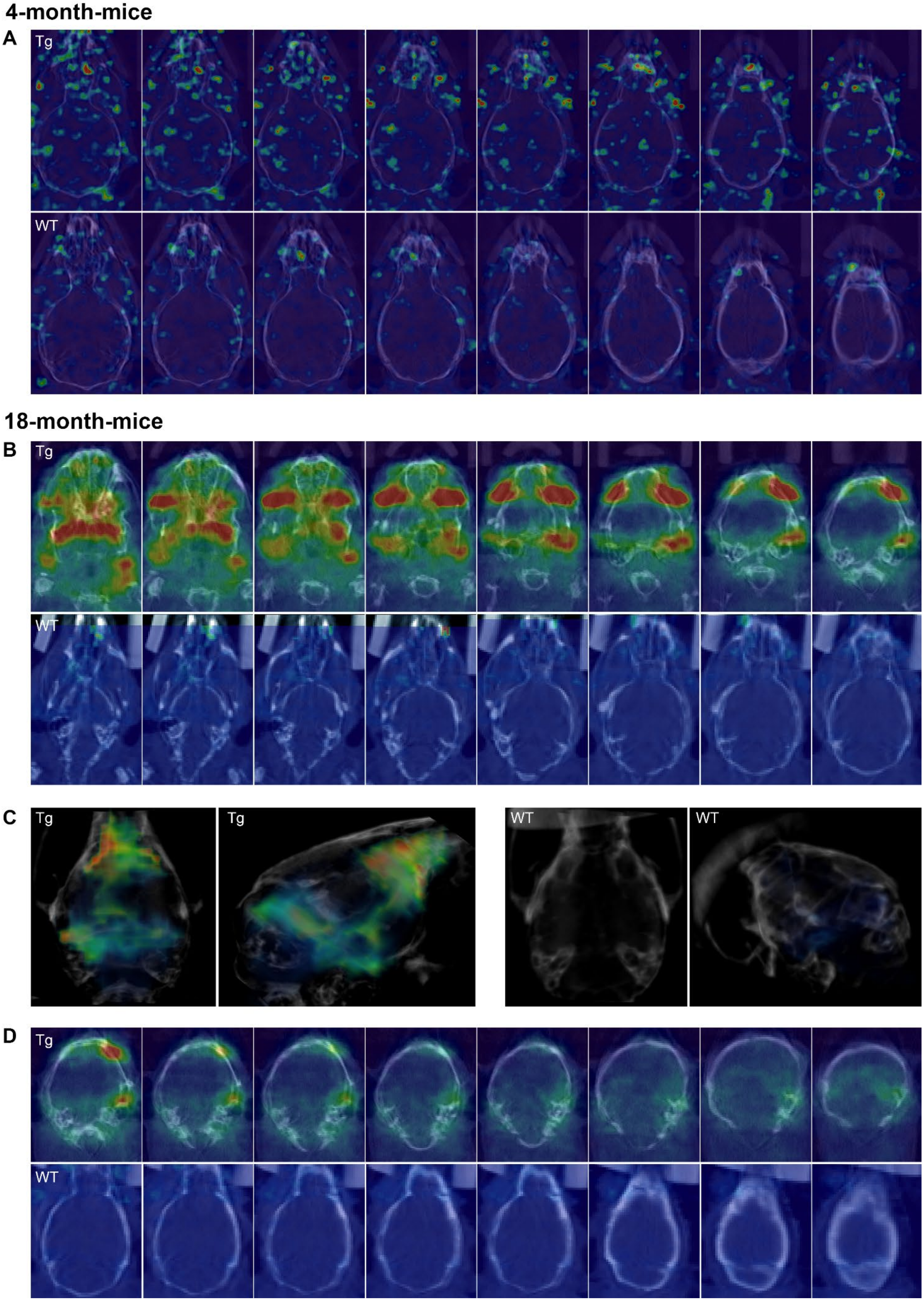
4-month-mice
A
Tg
WT
18-month-mice
B
Tg
WT
C
Tg
Tg
WT
WT
D
Tg
WT

### Supplemental Figure 4

**ACUPET probe gives 5xFAD selective PET signal in 4- and 18-month-old mice.**

Data were collected at approximately 4 hours after tail-vein injection for the 4-month-old mice (Panel A) and at approximately 1 hour after tail-vein injection for the 18-month-old mice (Panels B, C, and D). Panel A shows serial sections through the brains of 4-month-old Tg (top) and WT (bottom) animals. Some signal is detectable in the sinuses of both animals (indicating probe that is still circulating through the blood stream and has not yet fully accumulated in the liver and kidneys), but is significantly lower in the WT animal. Signal is also detected in the retina and brain of the Tg mouse, but no appreciable signal is detectable in the retina or brain of the WT animal.

Panels B-D show the serial sections and volume renderings for the 18-month-old Tg and WT mice. Due to the head of the Tg mouse being tilted forward 45 degrees, serial sections were collected through much of the nose and eyes before frontal cortex ($3^{rd}$ or $4^{th}$ image from the left of Panel B). All other animals were scanned with their heads flat in the holder.

Panel B shows a large signal in the retina. The high retinal signal in the Tg mouse is in agreement with the large volume of literature published in recent years showing that Aβ accumulates in the retina. In fact, the retina may prove to be an even earlier and/or more sensitive location for early detection of AD. Panel B also shows a large signal in the region of the carotid artery, near the shoulder/neck region, in the Tg mouse (based on views and analysis done using the mutiplanar view in Amira). Panel C shows 3D-volume rendering of the signal only in the brain. The skull was used as a boundary to calculate the signal within the brain and used to generate the volume renderings. Panel D shows the signal in serial sections starting from behind/beyond the eyes. There is signal detected in the frontal cortex and throughout the brain. The signal appears to be strongest in the frontal cortex, olfactory bulb, and entorhinal cortex/mid brain regions. Of note, AβO accumulation is also detectable in the fatty tissues of the Tg mice, both younger and old.

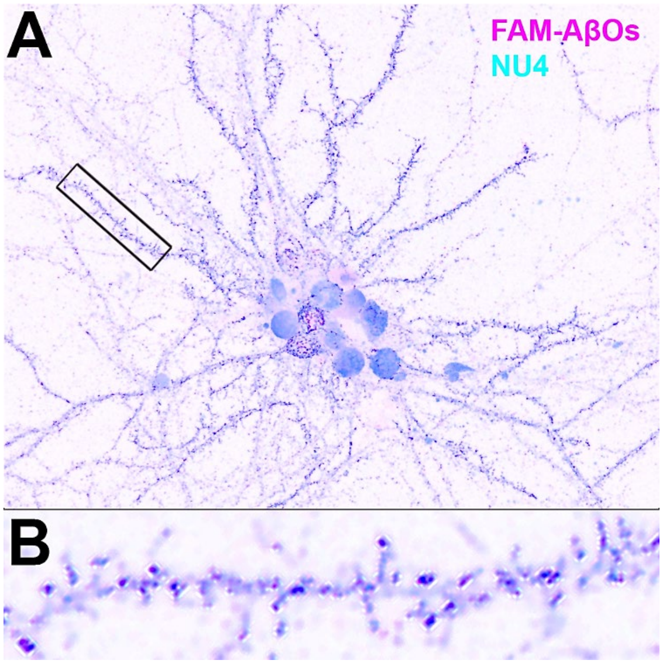


**Supplemental Figure 5**

**NU4-DOTA colocalizes with exogenous FAM-AβOs on cultured hippocampal neurons.**

NU4, covalently conjugated to DOTA (a chelator), is used as the basis for the AβO selective PET tracer. Primary rat hippocampal cells were incubated with fluorescent AβOs (FAM-AβOs; magenta) for 1 hour and then probed with NU4-DOTA followed by a mouse secondary (cyan). Data show high co-localization between AβOs and NU4-DOTA (dark blue).

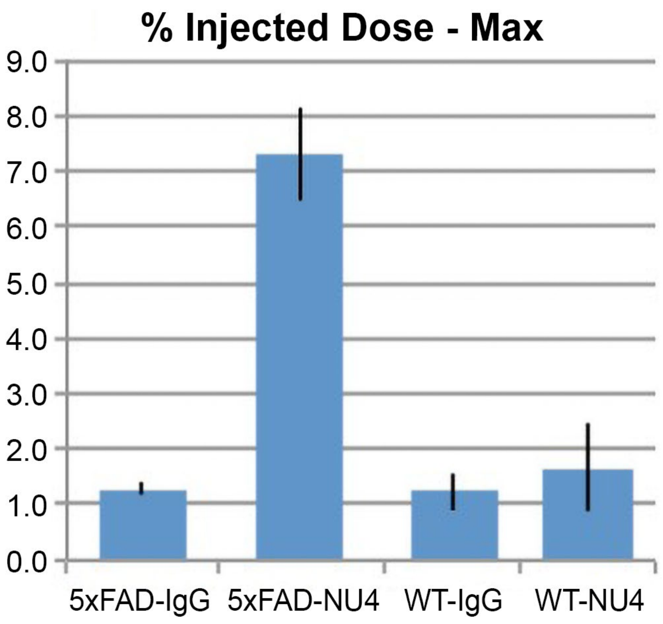


**Supplemental Figure 6**

**Specific uptake value (SUV) for NU4PET probe is significantly elevated for 5xFAD brain.**

Standard uptake values for NU4 PET were calculated using percentages of the injected dose/gram (%ID/g) as described in the methods. Data show that the signal detected by NU4PET in 5xFAD mice was significant in comparison to that detected in WT animals or 5xFAD animals injected with IgGPET.

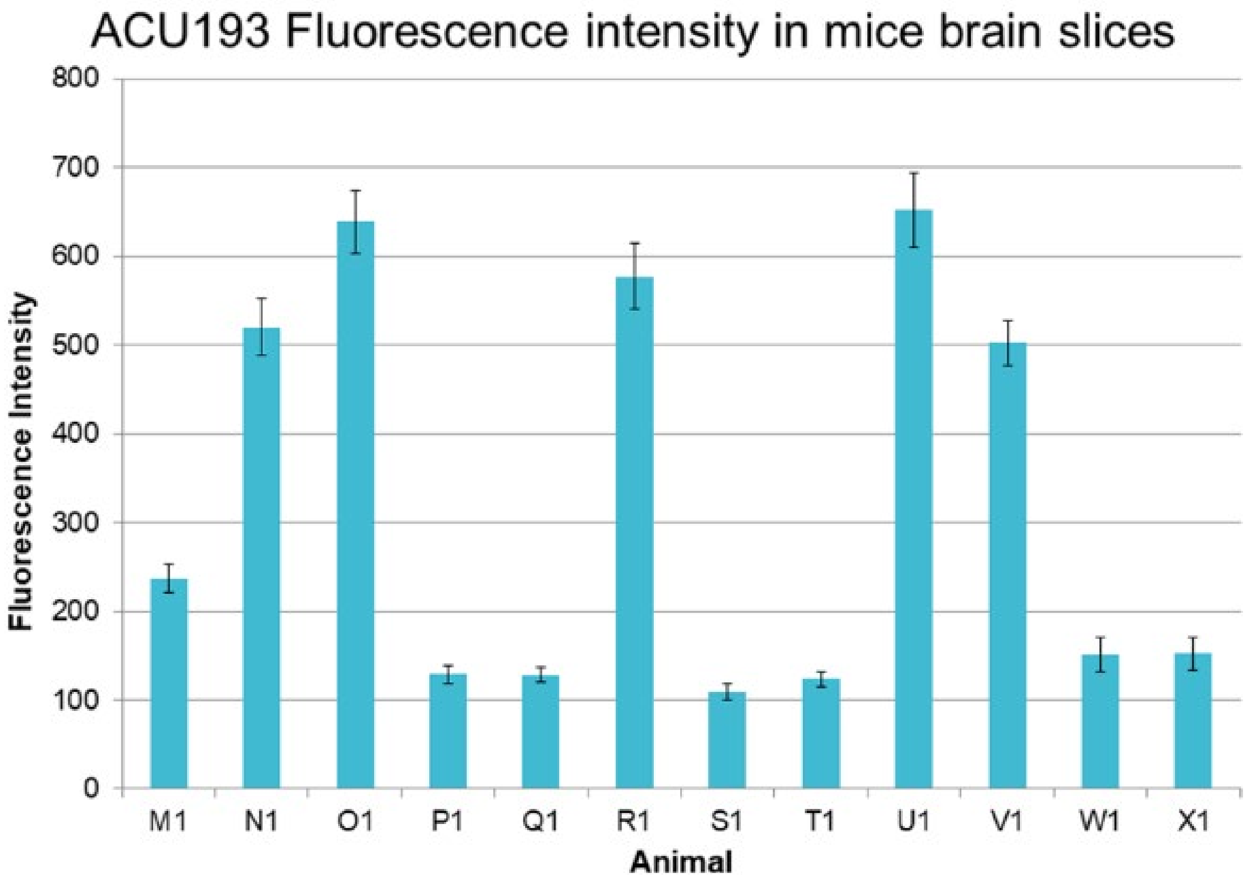


**Supplemental Figure 7**

**IHC confirms selective presence of AβOs in all 6 Tg imaged by PET.**

To confirm that oligomers were present in the Tg mice used for the PET experiments, mouse brains were fixed and stored until no longer radioactive. Blinded brain hemispheres were sagittally sliced and probed with ACU193 mAb against AβOs. A series of 25 images from the hippocampal and frontal cortex regions were collected. Using integrated densitometry, the average intensity was calculated and plotted for each brain. Data show that all six Tg mice (M1, N1, O1, R1, U1, V1) showed significant AβO signal over the WT animals (P1, Q1, S1, T1, W1, X1), although the signal in one Tg animal was half as intense as the other Tg mice

REFERENCES


Lacor, P.N., Buniel, M.C., Chang, L., Fernandez, S.J., Gong, Y., Viola, K.L., et al. (2004). Synaptic targeting by Alzheimer's-related amyloid beta oligomers. *J Neurosci* 24(45)**,** 10191-10200. doi: 10.1523/JNEUROSCI.3432-04.2004.

Lacor, P.N., Buniel, M.C., Furlow, P.W., Clemente, A.S., Velasco, P.T., Wood, M., et al. (2007). Abeta oligomer-induced aberrations in synapse composition, shape, and density provide a molecular basis for loss of connectivity in Alzheimer's disease. *J Neurosci* 27(4)**,** 796-807. doi: 10.1523/JNEUROSCI.3501-06.2007.

Pitt, J., Wilcox, K.C., Tortelli, V., Diniz, L.P., Oliveira, M.S., Dobbins, C., et al. (2017). Neuroprotective astrocyte-derived insulin/insulin-like growth factor 1 stimulates endocytic processing and extracellular release of neuron-bound Abeta oligomers. *Mol Biol Cell* 28(20)**,** 2623-2636. doi: 10.1091/mbc.E17-06-0416.
